# Matrix structure and microenvironment dynamics correlate with chemotherapy response in ovarian cancer

**DOI:** 10.1101/2025.05.19.654864

**Authors:** Florian Laforêts, Panoraia Kotantaki, Samar Elorbany, Joseph Hartlebury, Joash D Joy, Beatrice Malacrida, Rachel C Bryan-Ravenscroft, Chiara Berlato, Erica Di Federico, John F Marshall, Ranjit Manchanda, Wolfgang Jarolimek, Lara Perryman, Eleni Maniati, Frances R Balkwill

## Abstract

Elevated extracellular matrix (ECM) in the tumor microenvironment (TME) is associated with chemoresistance and poor prognosis. We hypothesized that modifying the ECM may enhance response to chemotherapy. We measured chemotherapy-induced changes in the TME of two mouse models of high-grade serous ovarian cancer (HGSOC) that differed in chemotherapy response. Treatment of the chemo-sensitive tumors triggered dynamic transcriptional ECM and immune changes and structural modifications of ECM proteins. These changes, observed over twenty-days post-chemotherapy, had relevance to HGSOC patient responses to chemotherapy. Integrating transcriptomics with ECM structure metrics, we identified ECM targets, including lysyl oxidase (LOX), that might enhance chemotherapy in less responsive mouse tumors. Given alone or in combination with chemotherapy, a pan-LOX inhibitor (PXS-5505) modulated fibroblast and immune cell distribution and reduced tumor stiffness in HGSOC chemo-resistant mouse tumors. Moreover, pre-treatment with PXS-5505 improved response to chemotherapy. We conclude that pretreatment with ECM targeting agents may improve response to chemotherapy, by altering ECM structure and immune responses.

## Introduction

High extracellular matrix (ECM) remodeling (fibrosis) and fibroblast/wound healing transcriptional signatures predict poor response to therapy and unfavorable prognosis in many human cancers (1–4). For instance, our previously published multi-level deconstruction of human omental metastases of high-grade serous ovarian cancer (HGSOC) revealed a pattern of extracellular matrix (ECM) molecules, the matrix index, that is associated with poor prognosis not only in this malignancy, but also in twelve other common cancer types(5).

The majority of HGSOC patients present with advanced disease that has spread throughout the peritoneum and omentum (6). They are generally treated with surgery and chemotherapy, often with three cycles of carboplatin/paclitaxel neoadjuvant chemotherapy (NACT) followed by debulking surgery, then a further three cycles of chemotherapy (7).

We previously reported that NACT induces anti-tumor immune activity in the human tumor microenvironment (TME) with effects on T cells, B cells and macrophages (8–12). However, these effects on innate and adaptive immune cells are not usually sufficient to induce long-lasting host anti-tumor activity. Most patients with HGSOC relapse within two years, with increasingly resistant disease (7).

In this paper, to further investigate the effects of chemotherapy on the TME, we have used two of our previously published syngeneic transplantable orthotopic mouse HGSOC models that replicate the human omental TME in terms of ECM composition and immune cells (13). The two intra-peritoneal mouse models varied significantly in their response to chemotherapy, with the 60577 model representing a highly chemo-responsive tumor and HGS2 representing a model with marginal response (13). The differences between the transcriptomes of untreated 60577 and HGS2 mouse omental tumors predicted response to chemotherapy in a HGSOC patient dataset, and significantly associated with ECM organization, cell adhesion, collagen catabolism and organization pathways(13). The weakly chemosensitive HGS2 model had higher levels of ECM proteins, increased numbers of fibroblasts and a greater matrix index than the chemosensitive 60577 model. Taken together, these results led us to propose that altering ECM composition and structure could enhance the effects of chemotherapy in a marginally responsive tumor.

To investigate this hypothesis, we conducted transcriptomic, spatial and structural analyses of the ECM in established omental tumors from our chemoresponsive model, 60577, after treatment with three cycles of carboplatin/paclitaxel chemotherapy. We related our results to patient samples and tested whether manipulating the ECM could enhance response in a less chemosensitive model, HGS2.

## Results

### Chemotherapy induces dynamic ECM and immune changes at the transcriptomic level in chemosensitive 60577 HGSOC murine tumors

We first studied the chemosensitive 60577 syngeneic peritoneal model of HGSOC, referred to above (13). The response of established tumors to three doses of carboplatin/paclitaxel (chemo) was confirmed (Figure 1A) showing a 12-week survival advantage of treatment. To replicate the NACT given to patients and relate to our previous studies of its effects on immune cells in the TME, we administered three doses of carboplatin/paclitaxel (chemo) to mice with established disease and collected tumors 24hours (TP1) and one week (TP2) after treatment. Tumors were also obtained three weeks after chemo (TP3), although insufficient mice in the control group were alive at this time. Omental tumors were weighed and divided for immunohistochemistry and bulk RNAseq (Figure 1B). There was a significant reduction of tumors burden at TP1 and TP2, which was maintained at TP3 (Figure 1C), in line with 60577 chemosensitivity seen in Figure 1A.

**Figure 1.**
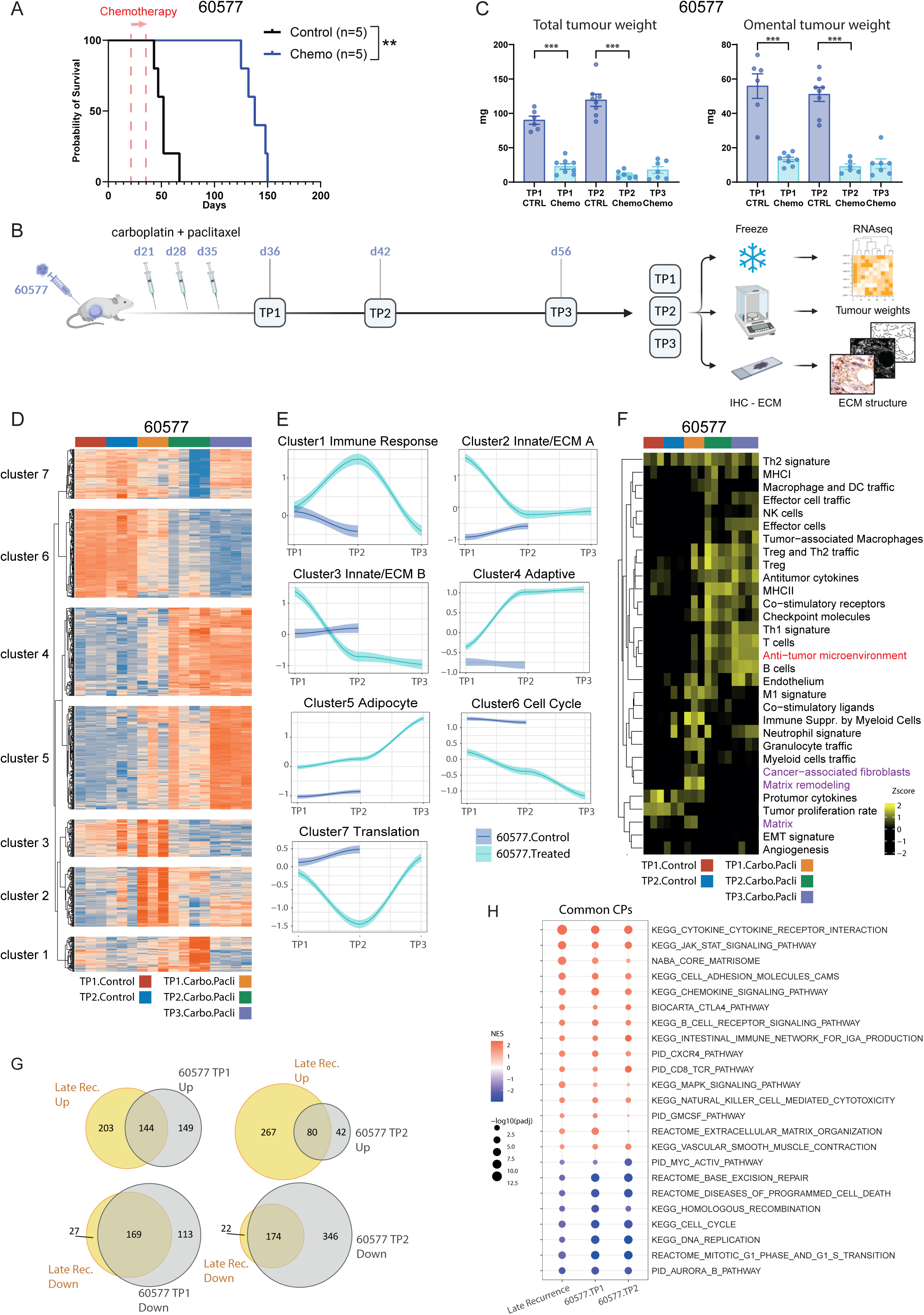
Response to chemotherapy associates with ECM and immune response pathway alterations in 60577 mouse HGSOC model. **A.** Response of 60577 tumor-bearing mice to three doses of a carboplatin and paclitaxel combination (Chemo, blue) compared to control group (black). Kaplan-Meier survival curve is shown, median survival for control and treated 60577 mice are 52 and 138 days, respectively. The log rank p value is depicted next to the survival curve. The start of the treatment is indicated by the red arrow. Numbers of mice enrolled in each arm are shown in parentheses. **B.** Scheme of animal study designed to assess the effect of carboplatin and paclitaxel on 60577 tumors, at different time points. Carboplatin (20 mg/kg) and paclitaxel (10mg/Kg) were administered i.p. once a week for three weeks starting 3 weeks post tumor cell injection. Mice were culled one day (TP1), one week (TP2) and three weeks (TP3) after last dose of chemotherapy, their tumors were resected, frozen or formalin-fixed and paraffin embedded after being weighted. RNA was extracted from the frozen tumors and subjected to RNAseq analysis. FFPE sections were stained for ECM markers to assess ECM structure and texture. **C.** Total and omental tumor weights from animals described in B. Each dot represents a tumor from an individual mouse, data represented as mean ±SEM. With blue and light blue showing tumors weight from respectively control (CTRL) and chemotherapy-treated (Chemo) mice. P values for Mann-Whitney U test are depicted on the bar plots for comparisons between control and treated mice. No significant difference between treated TP1, TP2 and TP3 was detected (Kruskal-Wallis test). * p<0.05, ** p<0.01, *** p<0.001. **D.** Heatmap illustrating log2RPKM gene expression for genes with a significant change between 60577 Treated vs Control (TP1 and/or TP2) and a significant change between at least one of the three treated timepoint contrasts (TP1 vs TP2, TP2 vs TP3 and TP1 vs TP3) (adjp < 0.05 and log2FC > |1|). Heatmap was clustered using K-means clustering into 7 clusters. **E.** Graphical illustration of the patterns of gene expression across the 7 clusters. Trendlines for the expression of each cluster were plotted using scaled gene expression and the median value of replicate mice. Graphs were plotted on ggplot with trendline using the loess smoothing method. **F**. Heatmap illustrating gsva enrichment scores of Bagaev et al^12^ functional gene expression signatures in 60577 control and treated tumors. The anti-tumor microenvironment signature (red) was calculated using the M1 signature, Th1 signature and anti-tumor cytokines signature genes as well as the B cell signature genes. ECM-related signature are highlighted in fuchsia. **G.** Venn diagrams illustrating the overlap of significantly enriched canonical pathways in the Adzibolosu et al. human HGSOC dataset^13^ of Late Recurrence post-treatment vs pre-treatment samples and canonical pathways significantly changing in 60577 Treated vs Control at TP1 and TP2, adjp <= 0.05. **H.** Enrichment dotplot illustrating some of the common canonical pathways in both the human and mouse data.

We next carried out transcriptomic analyses of the control and treated 60577 omental tumors at TP1, TP2 and TP3. Hierarchical cluster analysis of expressed genes showed treated 60577 tumors clustered separately from controls (Figure S1A). We compiled all genes with a significant differential expression in control versus treated (at TP1 and/or TP2) and at least between one of the treated time points (TP1 vs TP2, TP2 vs TP3, TP1 vs TP3). We then used K-means clustering and identified seven differential expression clusters (Figure 1D and Table S1A). We plotted the patterns of gene expression in these clusters (Figures 1E) and observed the changes at TP1, TP2 and TP3 in treated 60577 tumors compared to the controls.

Interestingly, pathways involving ECM organization, integrin-mediated signaling and innate immunity, while transiently increased at TP1, decreased at TP2, a change that was sustained at TP3 (clusters 2 and 3). Other immune response pathways were transiently elevated at TP2 (cluster 1). As innate immune pathways decreased (clusters 2 and 3), adaptive immune response signaling increased and was sustained at TP3, as shown by cluster 4 (Figure 1E and S1B, Table S1B). As expected, mitosis, cell division and cell cycle pathways were reduced in treated tumors compared to control and continuously decreased between TP1 and TP3 (cluster 6). Conversely, treated tumors presented elevated lipid and adipocyte-related pathways which increased between TP1 and TP3, as the tumor responds (cluster 5), along with a transient decrease in translation at TP2 (cluster 7).

In summary, RNAseq analysis showed dynamic transcriptional responses twenty days post-treatment in omental tumors that responded significantly to chemo. These were predominantly related to non-malignant cells in the TME. Initial ECM and innate immune gene expression declined as adaptive immune response expression increased. This was accompanied by decrease in pathways associated with cell proliferation and an increase in lipid synthesis gene expression in this adipocyte-rich TME.

In order to understand the relevance of these transcriptomic changes to human cancers, we referred to functional gene expression signatures defined in a pan-cancer human TME transcriptome analysis of 10,000 human tumors (14). These pan-cancer signatures captured the stromal and immune compartment of the TME, as well as anti-tumor and tumor-promoting processes. Selecting the mouse orthologues of the human genes in the signatures, we again found dynamic changes in ECM molecules, matrix remodeling and cancer-associated fibroblast pathways, along with TME cell signatures during the three weeks after chemo (Figure 1F, Table S2). Notably, TME changes observed in 60577 tumors were sustained at TP3. In line with the analysis in Figure 1D, we observed a transient increase in matrix remodeling and innate immune signatures at TP1 in treated tumors, including a transient increase in signatures of immature myeloid cells and cancer associated fibroblasts, suggesting these cell types are part of the remodeling process in responsive tumors. This initial increase in innate processes was followed by an increase in adaptive immune signatures, which became evident at TP2 and were sustained at TP3, including genes involved in Treg and Th2 cell trafficking. Thus, results from the mouse model transcriptional evolution are comparative to HGSOC patients treated with chemotherapy hence may provide insights for the disease in patients.

Of particular interest were changes in the “anti-tumor microenvironment” signature enrichment score (red highlight Figure 1F). The signature is adapted from a compilation of the M1 signature, Th1 signature and anti-tumor cytokines as described in Bagaev et al(14). We added B cells to this signature as they relate to good prognosis in ovarian cancer(15–18). Expression levels of this anti-tumor microenvironment score (anti-TME) increased after chemotherapy and were sustained at TP3 (Figure 1F). At TP2, the TME in responding tumors resembled the immune enriched-non-fibrotic profile described in Bagaev et al.

We conclude that anti-tumor immune and ECM responses were induced by chemotherapy. To further relate our findings in the mouse model to patient data, we analyzed a transcriptomic dataset of paired samples from twenty-four HGSOC patients pre and post carboplatin/paclitaxel NACT(19). This publication reported significant immune changes in post-treatment samples in patients with late disease recurrence compared to those who relapsed early (i.e. chemo-sensitive versus chemo-resistant tumors)(19). Using mouse orthologues and comparing the changes in treated versus control TP1 or TP2 60577 tumors with the post versus pre-chemo late recurrence patient samples (i.e., mouse versus human ‘responders’), we found significant overlap in up- and down-regulated pathways between mouse and human responding tumors (hypergeometric test p < 0.0001, Figure 1G). Again, ECM and immune pathways were significantly related to response in both mouse and human HGSOC tumors (Figure 1H, Table S3).

In summary, the dynamic and sustained changes in ECM and immune pathways observed after chemotherapy in 60577 tumors were relevant to transcriptional changes in human cancers. They reflected changes seen in NACT-treated HGSOC patients(19).

### Structural alteration of the ECM is a marker of response to chemotherapy in 60577 tumors

To measure effects of chemotherapy on ECM molecules, we performed IHC on the omental tumors studied in Figure 1C, measuring protein levels of three key ECM molecules we previously identified of prognostic significance in patient samples, fibronectin (FN1), versican (VCAN) and collagen 1A1 (COL1A1)(5). We also performed histochemistry for fibrillar collagens using Masson’s Trichrome (M3C). There was a transient reduction in versican positive staining area at TP2 in 60577 tumors, but no consistent changes for any of the others in control or treated tumors (Figure 2A). However, TWOMBLI(20) (structural) and QuPath (Haralick texture features)(21) analyses (278 metrics in total) revealed significant differences in the structure of FN1, VCAN and COL1A1 between control and treated 60577 tumors. M3C staining was mostly unaffected by chemotherapy at TP2 (Figure 2B). The radar plots showed that chemo caused a significant reduction in high density matrix (HDM) and curvature with significant increases in gap for the three ECM molecules at TP2. Haralick contrast and lacunarity increased, whereas box counting fractal dimension (BCFD) and total length decreased for two of the three ECM molecules. These ECM alterations were dynamic and varied between the different time points, with some that were sustained at TP3 for instance Col1A1 (Figure 2B, Figure S2A).

**Figure 2.**
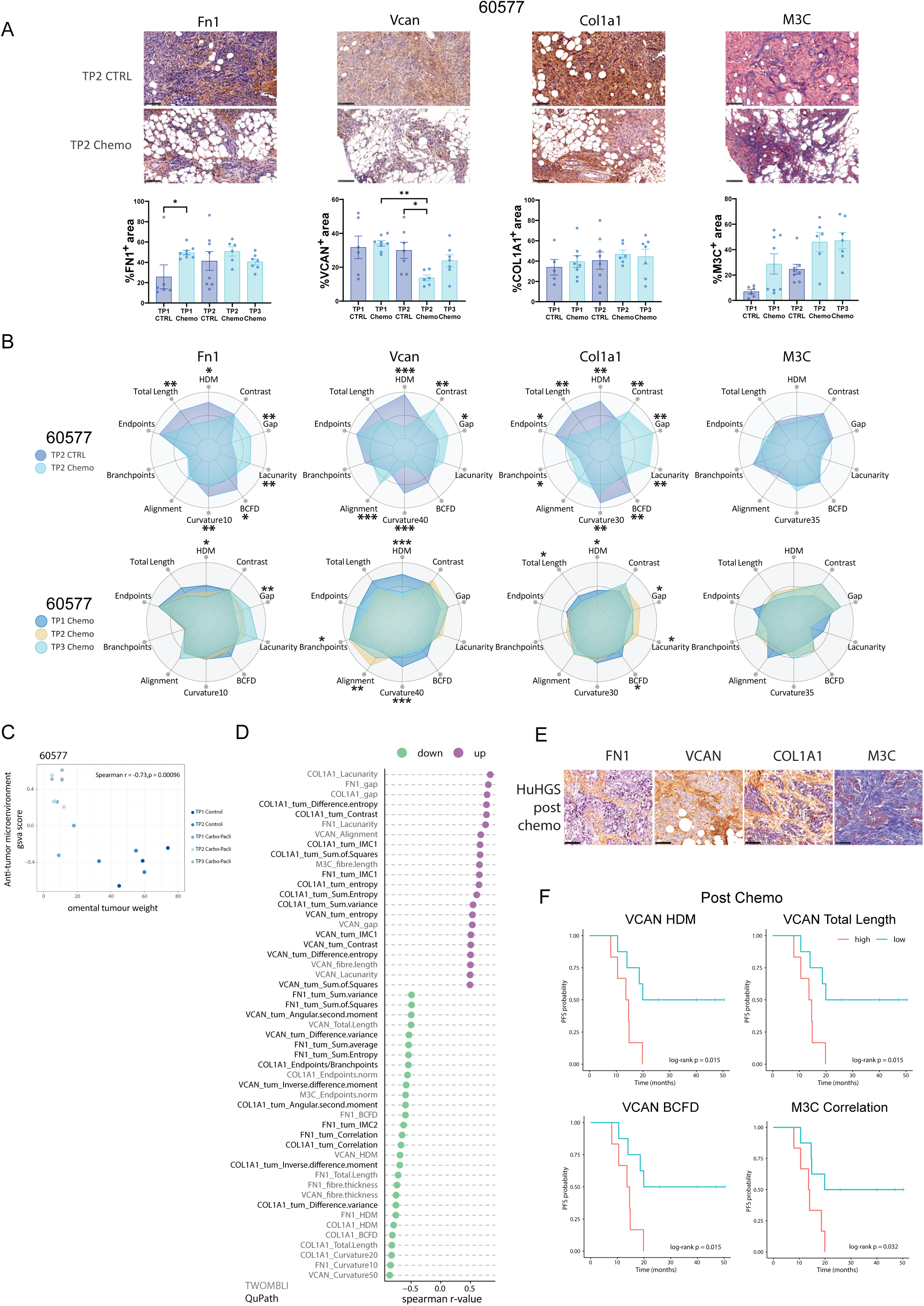
Structural alteration of the ECM is a marker of response to chemotherapy in 60577 tumors. **A.** Representative images of immunohistochemistry for fibronectin (FN1), versican (VCAN), Collagen 1A1 (COL1A1) immunohistochemistry and Masson’s Trichrome (M3C) histochemistry of tumors taken at TP2 and the corresponding quantifications of staining in 60577 (blue, control mice; light blue mice treated with carboplatin and paclitaxel) tumors at TP1, TP2 and TP3. Each dot represents one mouse. The Mann-Whitney test was used to determine statistical significance between control and treated groups for each individual time point. The difference between treated groups at time points 1, 2 and 3 (TP1, TP2 and TP3, respectively) was determined with the Kruskal-Wallis test’s multiple comparisons. **B.** Matrix patterns for FN1, VCAN, COL1A1 and Masson’s Trichrome (M3C) for 60577 tumors at TP2, assessed by TWOMBLI and Haralick feature analysis and visualised as radar plots. A selection of TWOMBLI metrics along with Haralick contrast is depicted. The average of structural and textural metrics was calculated for the samples analyzed in A. and the minimum and maximum range for each metric was computed across all categories. Values were normalized and scaled to the maximum range. Top panel, dark and light blue show the ECM pattern for control and treated TP2 tumors, respectively. Bottom panel shows ECM patterns for treated TP1, TP2 and TP3 tumors in blue, beige and cyan, respectively. HDM: High Density Matrix, Gap: Mean Gap Area, BCFD: Box Counting Fractal Dimension, Curvature10 to 40: Curvature metric assessed at windows of 10 to 40 pixels, Branchpoints: normalized branchpoints, Endpoints: normalized endpoints, Fiber Length: Average Fiber length. *p<=0.05, **p<= 0.01, ***p<= 0.001, Mann-Whitney test. **C.** Correlation scatter plot of the anti-tumor microenvironment scores with omental tumor weights in 60577. Each dot represents an individual mouse. Points that correspond to different experimental groups are illustrated in different colors. **D.** Cleveland plot showing spearman correlation coefficients of TWOMBLI and QuPath IHC image analysis metrics that correlated significantly with the anti-tumor microenvironment scores in 60577 (spearman p <= 0.05 and r >= |0.5|). **E.** Representative IHC images for FN1, VCAN, COL1A1 and Masson’s Trichrome staining (M3C) in human HGSOC omental biopsies. Scale bars, 100um. **F.** Kaplan-Meyer plots of progression-free survival on post-NACT samples; high and low groups were determined by median values nhigh = 7, nlow = 8.

### Structural ECM Features Associate with Anti-TME Transcriptomic Signatures

We next integrated these metrics with our transcriptomic data. We found a significant negative correlation between 60577 omental tumor weight and the anti-TME score described above (Figure 2C)(14), reinforcing the link between this signature and response to chemotherapy. We measured the correlation of all the ECM metrics in 60577 treated and control tumors across all timepoints with this enrichment score. Those that significantly correlated with the anti-TME score are displayed in a Cleveland plot (Figure 2D). This shows the quantitative, structural and textural ECM features that correlate positively (purple) and negatively (green) with the anti-TME score. Collectively, this gives a picture of decreasing HDM, total length, curvature, fiber thickness and increases in gap and lacunarity with the anti-TME score, supporting our hypothesis that ECM structure is involved in the response to chemotherapy.

In the absence of a reliable measure of response to chemotherapy at intermediary debulking surgery time point, we assessed if structural changes in ECM proteins associated with progression-free survival (PFS) in HGSOC patients, after NACT. We conducted the same IHC and histochemistry analysis in 15 HGSOC samples collected at interval debulking surgery 3-4 weeks after the third dose of NACT (Figure 2E). We looked for correlations between ECM structure and texture metrics with progression free survival (PFS). Several metrics correlated with progression-free survival in HGSOC patients, including VCAN Total Length and VCAN BCFD that were identified in the 60577 tumors as associated with response (Figure 2D), and M3C correlation (Figure 2F, Figure S2B).

Our results so far show that three doses of chemo trigger dynamic ECM and immune changes in the TME in a chemo-sensitive mouse HGSOC model. The ECM in chemo-treated 60577 tumors was less compact with increased gaps between fibers. Bioinformatic analysis showed relevance of these changes to HGSOC patient biopsies, and some ECM structural changes induced by chemo in the mouse tumors associated with progression-free survival in HGSOC patients.

### Chemotherapy fails to induce transcriptomic and ECM changes in a partially responsive murine HGSOC model, HGS2

Having established chemotherapy induces dynamic ECM and immune changes in the chemosensitive 60577 tumors, we asked if such changes were observed in a less responsive model. Treatment of established HGS2 tumors with six doses of chemo had limited survival benefit (Figure 3A) confirming our previously published results (22). We next treated HGS2 tumor-bearing mice with three doses of carboplatin and paclitaxel to resemble NACT (Figure 3B). As with the 60577 model, HGS2 control and chemo treated tumors were collected at the same three time points, TP1, TP2 and TP3. There was no HGS2 TP3 control group due to an insufficient number of control mice alive three weeks post chemo. Tumors were weighed and then prepared for histology and bulk RNAseq (Figure 3B). We observed no significant decrease in tumor weights at either time point (Figure 3C).

**Figure 3.**
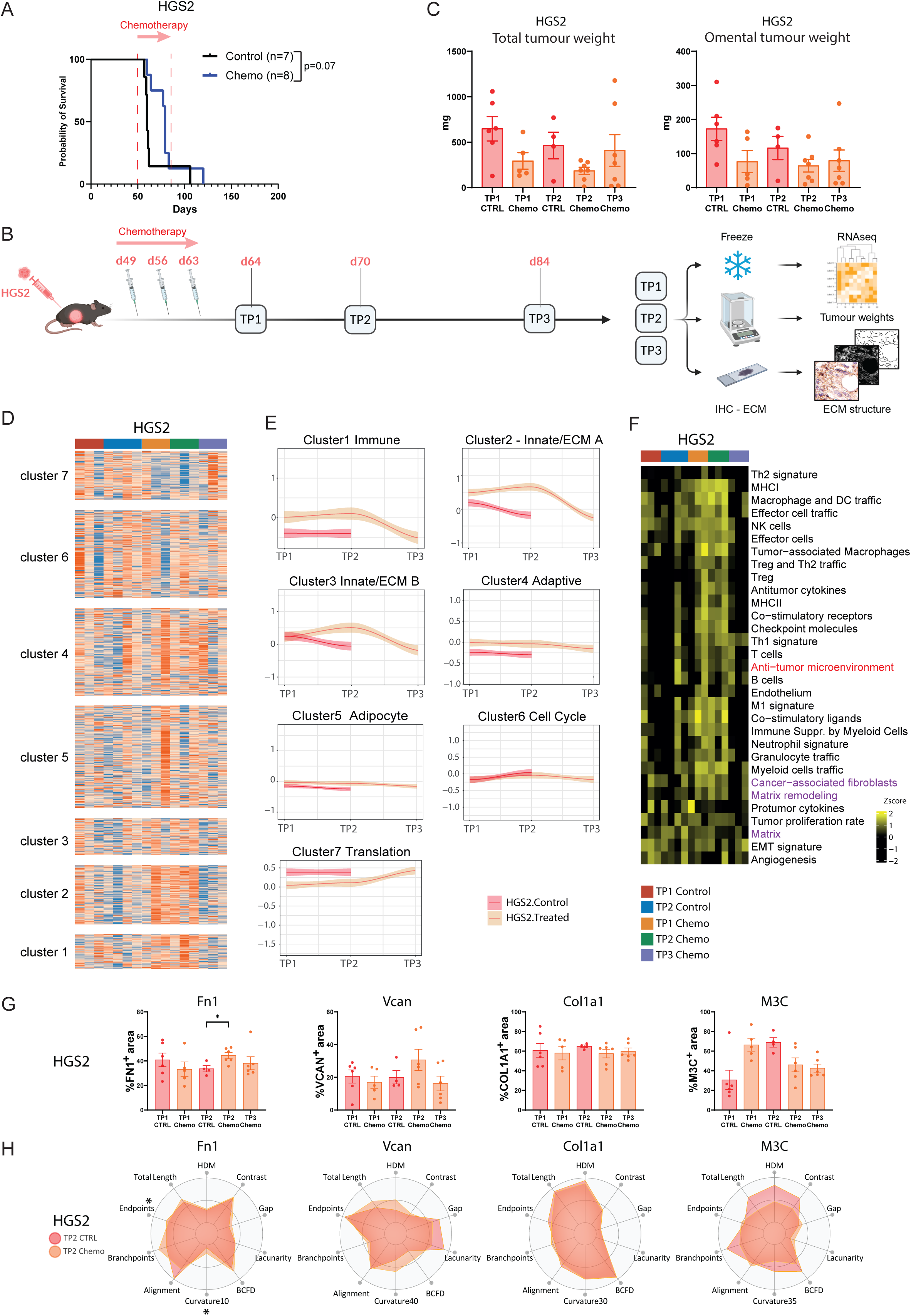
Chemotherapy fails to induce sustained transcriptomic and ECM changes in the HGSOC mouse model HGS2. **A.** Responses of HGS2 tumor-bearing mice to six doses of a carboplatin and paclitaxel combination (Chemo, blue) compared to control group (black). Kaplan-Meier survival curves are shown, median survival for control and treated HGS2 are 60 and 79 days, respectively. The log rank p value is depicted next to the survival curves. The start of the treatment is indicated by the red arrow. Numbers of mice enrolled in each arm are shown in parentheses. **B.** Scheme of animal study designed to assess the effect of carboplatin and paclitaxel on HGS2 orthotopic mouse model tumors, at time point. Carboplatin (20 mg/kg) and paclitaxel (10mg/Kg) were administered i.p. once a week for three weeks starting 7 weeks post tumor cell injection. Mice were culled one day (TP1), one week (TP2) and three weeks (TP3) after last dose of chemotherapy, their tumors were resected, frozen or formalin-fixed and paraffin embedded after being weighted. RNA was extracted from the frozen tumors and subjected to RNAseq analysis. FFPE sections were stained for ECM markers to assess ECM structure and texture. **C.** Total and omental tumor weights from animals described in **B**. Each dot represents a tumor from an individual mouse, data represented as mean ±SEM. With blue and dark orange are 60577 and HGS2 control mice shown respectively, whereas with light blue and orange are the respective treated samples from each mouse model colored. P values for Mann-Whitney U test are depicted on the bar plots for comparisons between control and treated mice. No significant difference between TP1, TP2 and TP3 was detected (Kruskal-Wallis test). **D.** Heatmap illustrating log2RPKM gene expression for genes making up the seven clusters identified in 60577 tumors in figure 1D, in HGS2 tumors. **E.** Graphical illustration of the patterns of gene expression across the 7 clusters in HGS2 tumors. Trendlines for the expression of each cluster were plotted using scaled gene expression and the median value of replicate mice. Graphs were plotted on ggplot with trendline using the loess smoothing method. **F**. Heatmap illustrating gsva enrichment scores from Bagaev et al.^12^ functional gene expression signatures in HGS2 tumor. The anti-tumor microenvironment signature was calculated using the M1 signature, Th1 signature and anti-tumor cytokines signature genes as well as the B cell signature genes. ECM-related signature are highlighted in fuchsia. **G.** Quantifications of fibronectin (FN1), versican (VCAN), Collagen 1A1 (COL1A1) immunohistochemistry and Masson’s Trichrome (M3C) histochemistry staining in HGS2 (red, control mice; orange, mice treated with carboplatin and paclitaxel) tumors. Each dot represents one mouse. The Mann-Whitney test was used to determine statistical significance between control and treated groups for each individual time point. The difference between treated groups at time points 1, 2 and 3 (TP1, TP2 and TP3, respectively) was determined with the Kruskal-Wallis test’s multiple comparisons. **H.** Matrix patterns for FN1, VCAN, COL1A1 and M3C for HGS2 tumors at TP2, as they were assessed by TWOMBLI and Haralick feature analysis and visualized in the radar plots. A selection of TWOMBLI metrics along with Haralick contrast is depicted. The average of structural and textural metrics was calculated for the samples analyzed in **B** and the minimum and maximum range for each metric was computed across all categories. Values were normalized and scaled to the maximum range. Red and orange show the ECM pattern for control and treated tumors, respectively. HDM: High Density Matrix, Gap: Mean Gap Area, BCFD: Box Counting Fractal Dimension, Curvature10 to 40: Curvature metric assessed at windows of 10 to 40 pixels, Branchpoints: normalized branchpoints, Endpoints: normalized endpoints, Fiber Length: Average Fiber length. *p<=0.05, **p<= 0.01, ***p<= 0.001, Mann-Whitney test.

Next, we conducted bulk RNAseq in HGS2 control and treated tumors at each time point. Hierarchical cluster analysis of expressed genes showed that, in this model, chemotherapy-treated tumors did not cluster separately from the controls (Figure S3A), and HGS2 omental tumor weight did not correlate with Bagaev et al’s anti-TME signature(14) (Figure S3B). We looked at expression of the genes that made up the seven clusters described in Figure 1D. These showed no significant differences in HGS2 (Figure 3D) between control and treated tumors, and plotting the cluster dynamic expression showed little to no variation over time (Figure 3E). We noted changes in HGS2 only using an unadjusted p value < 0.01. While we noted some overlap in differentially genes with 60577 (Figure S3C), looking at the genes that showed this low-level differential expression in control versus treated (at TP1 and/or TP2) in HGS2 tumors, we could not distinguish strong dynamic patterns of alterations between control and treated group (Figure S3D). Some genes with these low-level changes in treated compared to control groups at TP1 and TP2, were not sustained at TP3, suggesting a weak and transient response to chemotherapy (S3D).

We further used the pan-cancer human functional gene signatures identified by Bagaev et al to assess the relevance of these findings to human cancers. Looking at the mouse orthologues of those genes we found a limited number of differences between control and chemo-treated HGS2 tumors. Importantly, while some TME gene signatures showed weak alterations in treated tumors at TP1 and TP2, these reverted to control levels at TP3 (Figure 3F, Table S2).

We next assessed whether chemotherapy affected the ECM in HGS2 tumors. We stained for FN1, VCAN and COL1A1 by immunohistochemistry, and performed Masson’s Trichrome to assess fibrillar collagens (Figure 3G and S3E). Apart from a transient increase in FN1 at TP2, HGS2 tumors displayed stable levels of ECM molecules. Similarly, analysis of structure and texture features showed no difference between control and treated tumors at TP2 for any ECM marker analyzed (Figure 3H). Fibrillar collagen, as showed by Masson’s trichrome, presented strong alterations at TP1, with collagens overall denser, more complex and with less aligned fibers. This was only observed at TP1, however and was not sustained at TP2 and TP3 (Figure S3F and S3G). Together, our results with HGS2 tumor-bearing mice show this model responds weakly and transiently to chemotherapy.

### Integration of RNAseq profiles with ECM structure metrics reveals strategy to target HGSOC tumors with reduced chemosensitivity

Although, as shown above, 60577 and HGS2-tumor bearing mice displayed different sensitivity to chemotherapy *in vivo*, 60577 and HGS2 cells were equally sensitive to carboplatin or paclitaxel *in vitro* (Figure 4A) and displayed similar growth rates as monocultures (Figure 4B). This suggested that any difference in chemosensitivity was not simply due to an intrinsic differential sensitivity of the cells to chemotherapy but was affected by their engagement with, and modification of, the microenvironment as they grew in the peritoneum and omentum. Taken with our previously published data, this suggested that the lack of effect of the ECM structure in HGS2 is, in part, responsible for its limited chemosensitivity.

**Figure 4.**
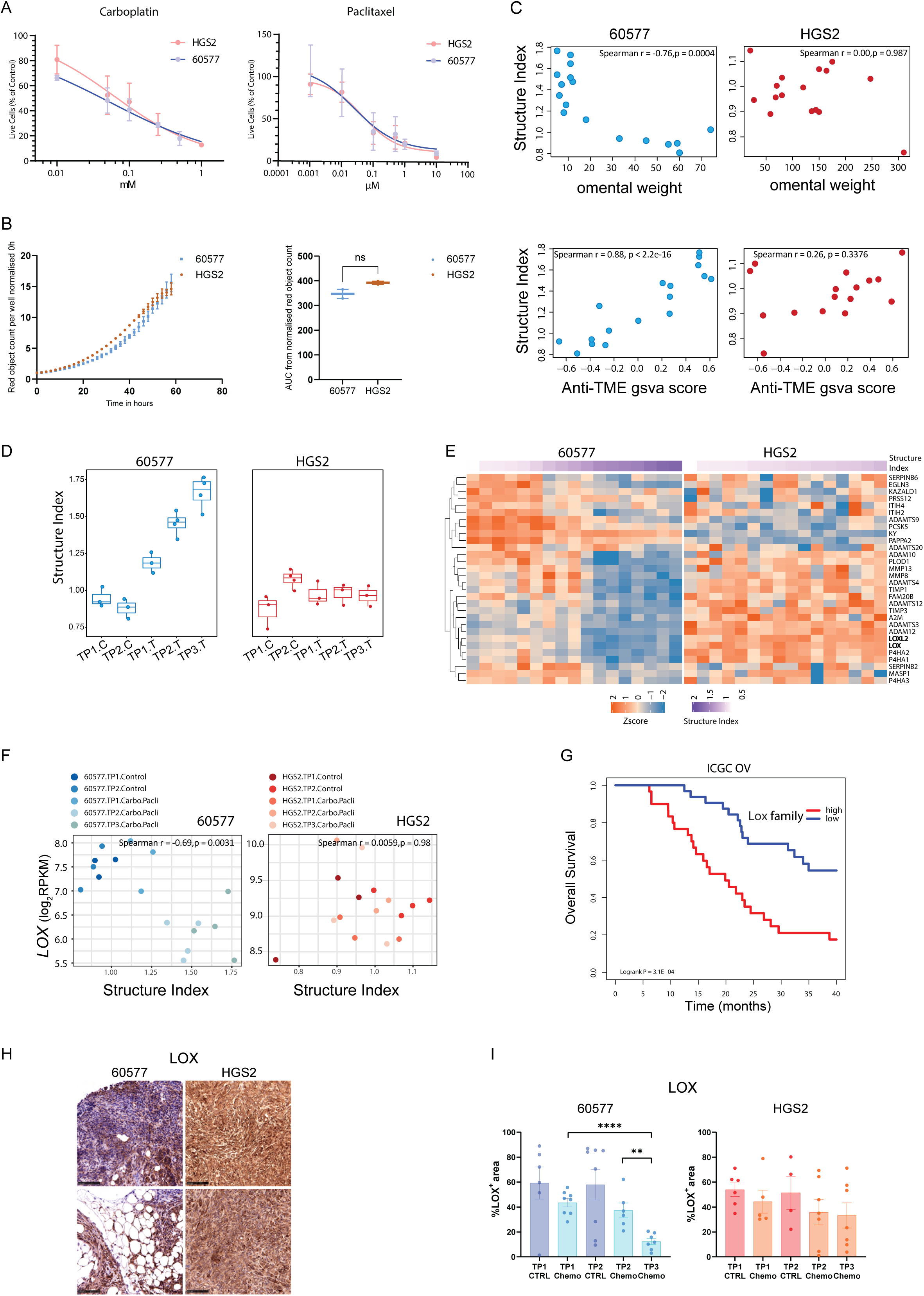
Integration of RNAseq profiles with IHC metrics uncovers therapeutic targets in chemoresistant HGSOC. **A**. Chemosensitivity of 60577 and HGS2 mouse cell lines to carboplatin and paclitaxel. The ratio of live cells treated with carboplatin and paclitaxel / live cells from control is shown. The results were plotted as mean ± standard error (SEM) of three separate experiments for each experimental condition. IC50 was tested with Students’ t-test and was not significant**. B**. Incucyte S3 curves showing the proliferation of HGS2 (red) and 60577 (green) cell lines (left) and area under the curve (AUC) of the curves on the right. Red lentivirus labelled cancer cells were plated (n=2 per cell line) and the plate was imaged every 2h in the incucyte S3. Error bars represent mean and SD. AUC of the proliferation curves on the left (n=2 replicates). Error bars at mean and SD. Two-sided unpaired student t-test. **C.** Correlation scatter plots illustrating association of Structure Index with omental tumor weight (top row) and anti-TME gsva score (bottom row) in 60577 (blue) and HGS2 (red). Each dot represents an individual mouse. The structure Index was calculated using the average of scaled IHC metrics with positive correlation with anti-tumor microenvironment gsva scores divided by the average of scaled metrics with negative correlation with anti-tumor microenvironment scores. **D**. Boxplots illustrate the structure index values in the 5 experimental groups for 60577 (blue) and HGS2 (red). Each dot represents an individual mouse **E**. Heatmap of gene expression of matrisome regulators with positive association with Structure Index in 60577 (Spearman’s r >= 0.5 and p <= 0.05) but not in HGS2 (p > 0.05). **F**. Correlation scatter plots illustrating *Lox* correlation with structure index in each mouse model, with each dot representing an individual mouse. **G.** LOX family gene expression gsva enrichment scores were calculated in the ICGC-OV dataset. Kaplan-Meyer plot of overall survival of Lox family high and low samples on ICGC-OV; high and low groups correspond to the top 33% and lower 33% samples ranked by decreasing lox family expression score nhigh = 31, nlow = 32. **H.** Immunohistochemical staining and **I.** Quantification for LOX in control and chemotherapy treated tumors from 60577 (indicated with blue) and HGS2 models (indicated with orange). % positive area in the tumor plus stromal area (excluding fat) was quantified in both cases. Each dot represents a mouse, p-values correspond to Kruskal-Wallis test and Mann-Whitney for individual time points * p<0.05, ** p<0.01, *** p<0.001. A representative image from TP2 is shown from each tumor, scale bar: 100um.

To identify ECM targets of potential importance to chemo response, we calculated a score, combining the ECM metrics that correlated with the anti-TME signature and tumor weight (Figure 2C, 2D). We termed this the ‘structure index.’ The structure index reflects features of the ECM that are associated with response to chemotherapy and can be computed for each sample. In 60577, a high structure index denoted an ECM structure linked to good response to chemotherapy, as shown by its positive correlation with the anti-TME gsva score, and negative correlation with omental tumor weight (Figure 4C), while a low structure index associated with poor response. This correlation was not observed in HGS2 tumors. The structure index increased with time after treatment in 60577 tumors, but did not change significantly in HGS2 (Figure 4D).

This gave us a score that we could integrate with transcriptomic data in each tumor. We focused on matrisome genes. Genes correlated with the structure index therefore also correlate with an ECM structure and response to chemo. 198 and 160 matrisome genes were respectively associated with a significant increase and decrease in structure index with chemo in 60577 and these included all six different categories of ECM genes (Figure S4A). We constructed heatmaps of those genes, selecting those that significantly decreased with chemotherapy in 60577 (as the structure index increased), but were largely unchanged in HGS2 (Table S4).

Because expression of individual genes in these heatmaps correlated with ECM structure and response to chemo, we reasoned this analysis might identify genes whose protein products could be targeted to generate ECM changes and improve response to chemotherapy. We focused on the ECM regulatory genes as potential targets for proof-of-concept (Figure 4E), selecting lysyl oxidase (LOX) and LOXL2. LOXs are a family of secreted copper-dependent enzymes with important roles in cross-linking of fibrillar collagen into mature ECM(23, 24). Untreated tumor gene expression levels of LOX and LOXL2 were lower in 60577 than HGS2 and showed a significant negative correlation with the structure index at TPs 1,2 and 3 in treated 60577 tumors but not HGS2, where levels remained high (Figure 4F and S4B). In addition, high expression of LOX family genes correlated with lower overall survival in the ICGC ovarian cancer database(25) (Figure 4G). There was a significant decline in LOX and LOXL2 in response to chemo in 60577 tumors but not HGS2 (Figure 4H, 4I, S4C and S4D). LOX was strongly linked to the regulation of ECM genes in a matrisome-based correlation network (Figure S4E). Moreover, in 60577 tumors, chemotherapy induced COL1A1 structure changes that were sustained at TP3 (Figure 2B). As we did not observe such alterations in HGS2, we hypothesized that targeting collagen cross-linking by inhibiting LOX in HGS2, would enhance the response to chemotherapy.

### Early exposure to PXS-5505 increases survival in HGS2 tumor-bearing mice when combined with chemotherapy

We tested our hypothesis using a recently described ECM modulating pan-LOX inhibitor PXS-5505(26) that inhibits *de novo* collagen cross-linking. We first treated tumors at an early stage of development, to assess if preventing build-up of excessive collagen crosslinking would enhance the effects of chemotherapy. We started oral treatment with the LOX inhibitor in chow seven days after HGS2 cell injection, commencing chemotherapy six weeks later (Figure 5A). Mice received six doses of chemo, to replicate the treatment regime of most HGSOC patients (7). Combining results from three experiments, we show that six cycles of chemotherapy resulted in a 2 week increase in mouse median survival (73 versus 89 days) (Figure 5B). ‘Early’ treatment with PXS-5505 alone had a small but significant effect on mouse survival (73 versus 81 days, p=0.002), with three long-term survivors (two at 203 days, one at 405 days) (Figure 5B). Early PXS-5505 treatment also significantly enhanced the effects of chemotherapy, although the improvement was modest (89 versus 98 days p=0.05).

**Figure 5.**
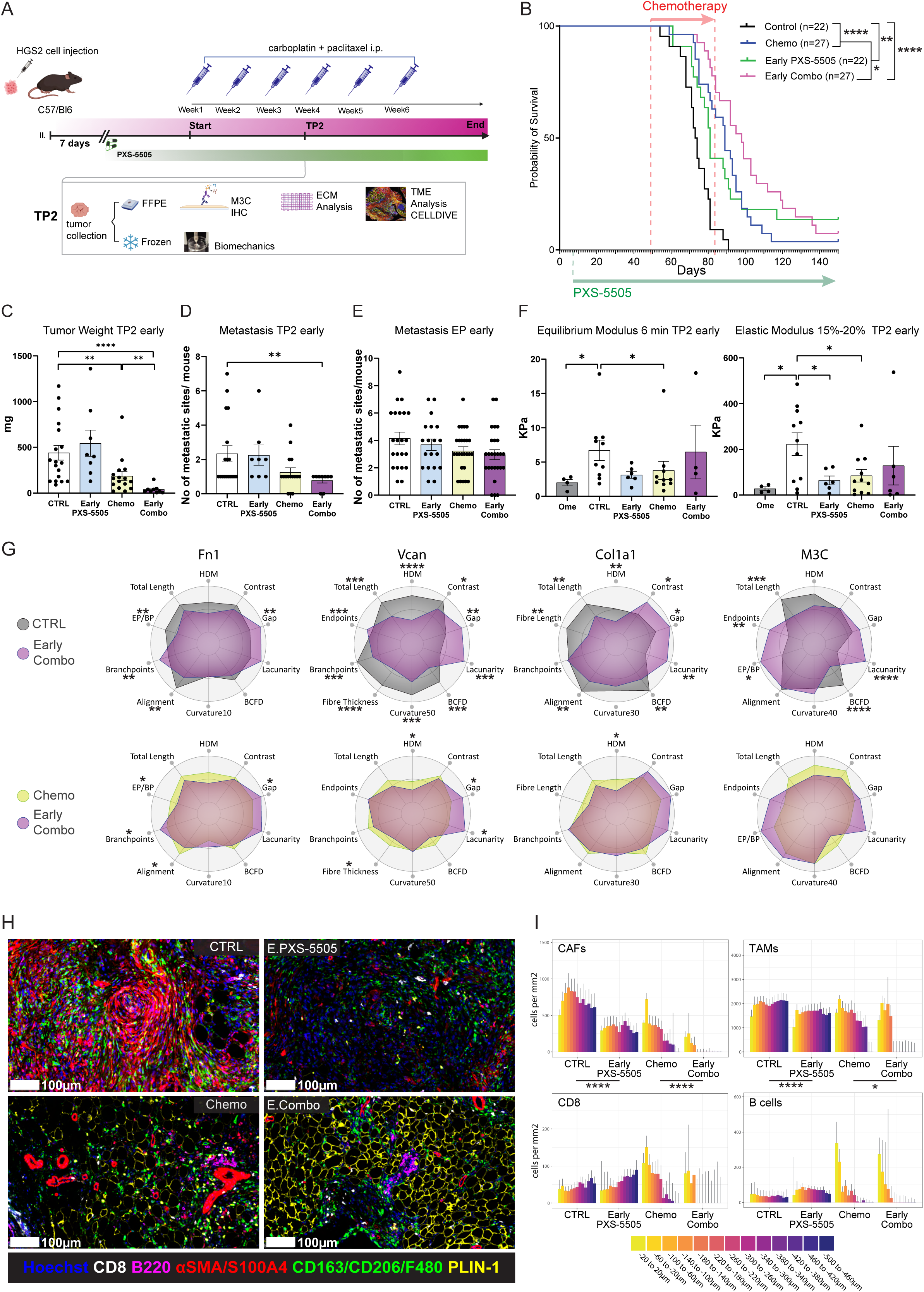
Prolonged exposure to PXS-5505 increases survival in HGS2 tumor-bearing mice, in combination with chemotherapy. **A**. Schematic diagram of *in vivo* HGS2 survival and time point (TP2) study and experimental outcomes. C57/Bl6 mice were injected i.p. with HGS2 cells and treated with 20 mg/kg carboplatin and 10mg/kg paclitaxel or vehicle once per week for three (TP2) or six cycles (endpoint). Onset of chemotherapy treatment commenced 49 days post tumor cell injection, whereas onset of PXS-5505 treatment commenced *ad libitum* 7 days post tumor cell injection (early treatment), where we have found colonization of the omentum by HGS2 tumor cells and continued to endpoint. Estimated dose of PXS-5505 *ad libitum* was 170mg/kg LOXi which is equivalent with a 10mg/kg oral gavage. **B.** Response of mice injected with HGS2 to six cycles of chemotherapy and/or to early treatment with PXS-5505 *ad libitum*. Median survival for control 73 days, for chemo 89 days, early PXS-5505 81 days, early Combo 98 days. The log rank p value is depicted on the survival curves. The start of the treatment indicated by the red arrow for chemotherapy, the green arrow for PXS-5505. Number of mice in each arm in parentheses, data pooled from three independent experiments. **C-E**. Quantification of total tumor weight, metastasis analysis at TP2 and endpoint (EP) across all treatment groups, following treatment plan depicted in 5A. Data are derived from one experiment combined with data from control and chemotherapy treated mice from two additional independent experiments. For weights, data are shown as mean ± sem and p-values correspond to two-tailed Mann-Whitney U test. Overt metastatic lesions in different anatomical regions were recorded with each dot representing one mouse, control= 18, chemo=16, early PXS-5505=8 and early combo=9. **F.** Biomechanical assessment of control and early-treated HGS2 tumors, at TP2. Equilibrium modulus as well as elastic modulus within the range of 15–20%, were calculated for TP2 across all treatment groups following treatment plan as depicted in **A**. Normal omenta (18 weeks) n=4, control n=10-11, chemo n=10-11, carly PXS-5505 n=6, early combo n=4-6. P-values correspond to two-tailed Mann-Whitney U test. **G.** Structural and textural modifications of ECM induced by early combination treatment with PXS-5505 and chemo in HGS2 tumors at TP2 compared to control and chemo. Radar plots show selected metrics for FN1, VCAN, COL1A1 and M3C fibers. The scaled average for each group is depicted. Control, grey, early combo, purple, chemo, yellow. P-values correspond to two-tailed Mann-Whitney U test. Data pooled from two individual experiments, along with controls and chemo treated samples from a third experiment. Control n=16-18, chemo n=16, early Combo n=9. **H.** Representative images of tumors from each treatment group stained for tumor-associated macrophages (TAMs, CD163/CD206/F480, green), adipocytes PLIN1 (yellow), CD8 (white), B220 (fuchsia) and cancer-associated fibroblasts (CAFs, αSMA/S100A4, red). Hoechst 33342 is in blue; scale bar is at 100um. **I.** The distribution of CAFs, TAMS, CD8 and B cells was recorded using the infiltration analysis tool from HALO in 40um wide zones from malignant cell areas in all four treatment groups, over a 500um range. p-values correspond to two-way ANOVA of cell density comparing treatment groups, taking zones into account. * p<0.05, ** p<0.01, *** p<0.001.

We analyzed tumors one week after the third dose of chemo, equivalent to TP2, with these treatment combinations. We chose TP2 as this time point displayed the strongest ECM structure changes in 60577 chemo treated tumor. Early PXS-5505 treatment enhanced the tumor weight reduction caused by chemo (Figure 5C) and the number of metastases was significantly reduced in the combination group compared to control (Figure 5D). When mice reached end point, there was no difference in numbers of metastases (Figure 5E).

We evaluated tumor stiffness using stress relaxation tests with unidirectional indentation, calculating both the elastic modulus at 15–20% strain and the equilibrium modulus. The elastic modulus reflects tissue stiffness under moderate, physiologically relevant deformation, providing insights into ECM structure, disease state, and treatment response(27). Higher values often indicate fibrosis or matrix cross-linking, poor drug penetration, or immune exclusion(28), while lower values suggest ECM softening or remodeling(29). Higher values of equilibrium modulus indicate persistent matrix stiffness, while lower values suggest degradation or therapeutic remodeling(27, 28). Elastic modulus was significantly reduced with early PXS-5505 treatment and with chemo (Figure 5F). We stained for FN1, VCAN, COL1A1 and M3C. We quantified these staining and analyzed their structure and texture. There were no changes in protein density apart from VCAN which decreased with chemo, a change that was sustained in the combination treatment (Figure S5A). Early PXS-5505 treatment alone altered M3C and FN1 structure compared to control, further altered VCAN and reduced COL1A1 HDM in the combination treatment compared to chemotherapy (Figures 5G and S5B). It was interesting to note some changes to the ECM in the chemo-treated HGS2 tumors had similarities to the 60577 responses described earlier (Figures 2B, and S5B), supporting our conclusion that chemotherapy affects the ECM, albeit less so in HGS2 tumors than 60577.

As chemo had also altered the transcriptional patterns of immune cells in 60577, we used spatial multiplex IHC to study the HGS2 TME with a total of seventeen different markers. Representative images with a selection of markers are shown in Figures 5H and S5C. Measuring the density of TME cells in the 500μm region from the outer edge towards the malignant cell area, we noted a significant reduction in the density of cancer-associated fibroblasts (CAFs, aSMA+ and/or S100A4+/- and CK14 and 19 and F4/80) and tumor-associated macrophages (TAMs, CD206+ and/or CD163+ and F4/80+ and aSMA-) with early PXS-5505, both as monotherapy and in combination with chemotherapy (Figure 5I), while B and CD8^+^ T cell numbers and distribution remained unchanged.

In conclusion, if given early during tumor development, PXS-5505 alone altered the TME of established tumors, reducing stiffness and opening-up the space between ECM fibers. It also reduced the distribution and density of fibroblasts and TAMs and enhanced survival when combined with chemotherapy.

### LOX inhibition in established tumors transiently decreases stiffness, alters ECM structure and the immune infiltrate but does not deliver a sustained response when combined to chemotherapy

To mimic combination of PXS-5505 with chemotherapy in a more clinically-relevant setting, we assessed whether PXS-5505 treatment of established tumors, commencing concomitantly with chemotherapy seven weeks after HGS2 cell injection, also enhanced the effect of chemotherapy (Figure 6A). We combined results from three separate experiments, two of which had both adjuvant (early) and concomitant (late) PXS-5505 arms, therefore the control and chemotherapy arms are the same as Figure 5B. We measured a decrease in LOX activity in the aortas of PXS-5505-treated mice, confirming that drug delivery in chow was effective (Figure S6A). Individual experiments from Figs. 5 and 6 are shown in Figure S6B. While PXS-5505 monotherapy did not affect median survival, there was a small improvement when it was given with chemotherapy compared to chemotherapy alone (95 days versus 89) (p=0.065).

**Figure 6.**
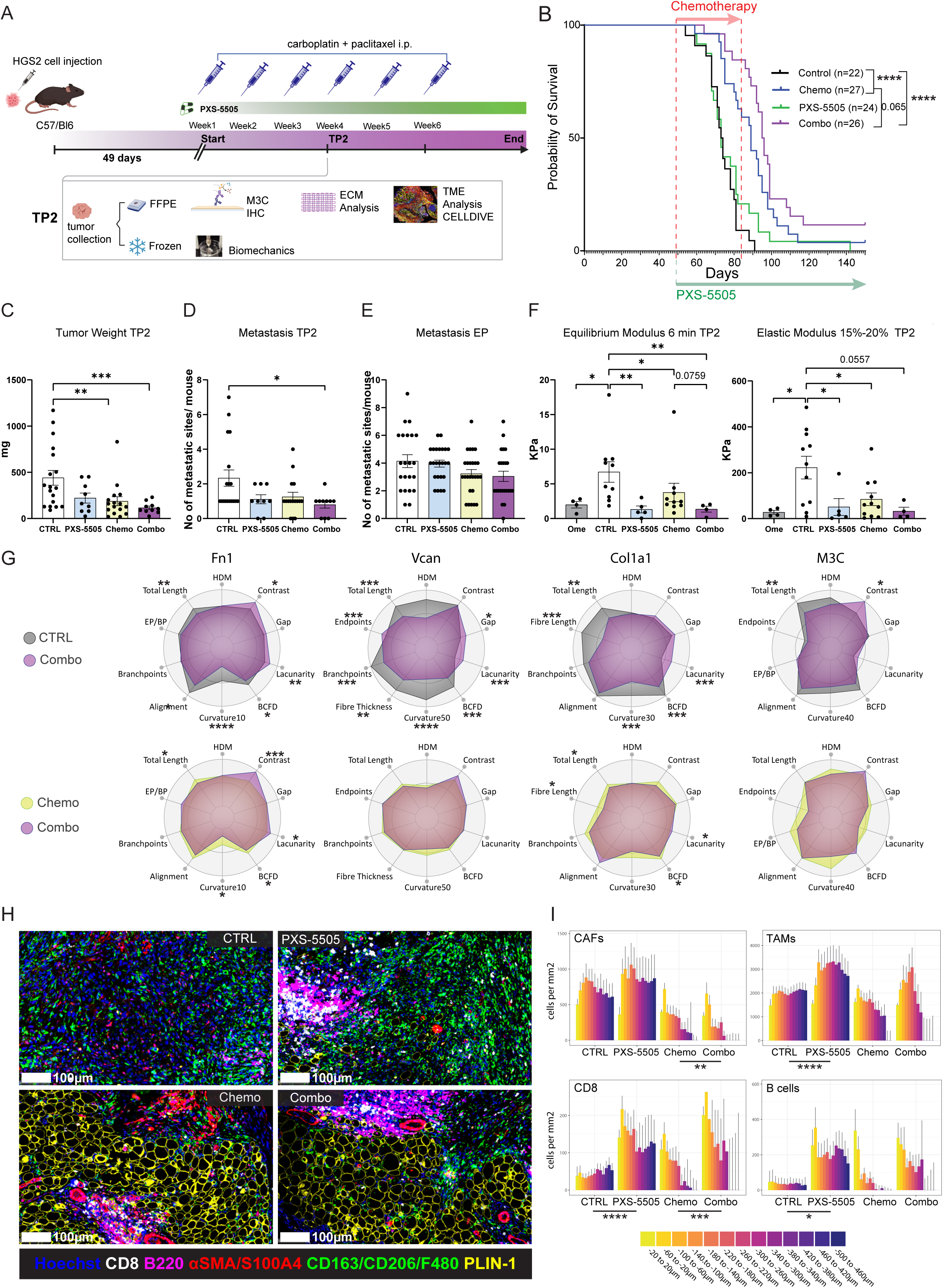
Concomitant chemotherapy and LOX inhibition decreases tumor stiffness, alters ECM structure and the immune infiltrate. **A**. Schematic diagram of *in vivo* HGS2 survival and time point (TP2) study. C57/Bl6 mice were injected i.p. with HGS2 cells and treated with 20 mg/kg carboplatin and 10mg/kg paclitaxel once per week or vehicle for three (TP2) or six cycles (endpoint) and/or PXS-5505. Onset of chemotherapy (i.p.) and PXS-5505 (*ad libitum*) treatment commenced with first detection of palpable tumors (49 days post tumor cell injection). Estimated dose of PXS-5505 *ad libitum* was 170mg/kg LOXi which is equivalent with a 10mg/kg oral gavage. M3C= Masson’s Trichrome. **B.** Response of mice injected with HGS2 cells to six cycles of chemotherapy and/or treatment with PXS-5505 *ad libitum*. Survival curve is shown and median survival for control is 73days, chemo 89days, late PXS-5505 73days, combo 95 days. The log rank p value is depicted on the survival curves. The start of the treatment is indicated by the red arrow for chemotherapy, green arrow for PXS-5505. Number of mice enrolled in each arm shown in parentheses, data pooled from three independent experiments. **C-E**. Quantification of total tumor weight at TP2, metastasis analysis for TP2 and endpoint (EP), respectively. Overt metastatic lesions in different anatomical regions were recorded, with each dot representing one mouse. Control= 18, Chemo=16, Late PXS-5505=9 and Combo=10. Data derived from two independent experiments combined with data from control and chemotherapy treated mice from a third experiment. For weights, data are shown as mean ± sem. p-values correspond to two-tailed Mann-Whitney U test. **F.** Biomechanical assessment of control and treated HGS2 tumors. Equilibrium modulus, as well as elastic modulus within the range of 15–20%, were calculated for TP2 across all treatment groups following treatment plan as depicted in **A.** For TP2: Normal omentum (OmeEach dot represents a tumor or tissue from an individual mouse. p-values correspond to two-tailed Mann-Whitney U test. **G**. Structural and textural modifications of ECM induced by late combination treatment with PXS-5505 and chemo in HGS2 tumors at TP2 compared to control and chemo. Radar plots computed as previously, show selected metrics for FN1, VCAN, COL1A1 and M3C fibers. The scaled average for each group is depicted. Control in grey, late combo in purple, chemo group in yellow. P-values correspond to two-tailed Mann-Whitney U test. Data pooled from two individual experiments are shown, along with controls and chemo treated samples from a third experiment. Control n=16-18, Chemo n=16, Late PXS-5505 n=8, late combo n=10. **H.** Representative image panel of a tumor from each treatment group stained for tumour-associated macrophages (TAMs, CD163/CD206/F480, green), PLIN1 (yellow), CD8 (white) and B220 (fuchsia) and cancer-associated fibroblasts (CAFs, αSMA/S100A4, red). Hoechst 33342 is in blue; scale bar is at 100um. **I.** The distribution of CAFs, TAMS, CD8 and B cells was recorded using the infiltration analysis tool from HALO in 40um wide zones from malignant cell areas in all four treatment groups, over a 500um range. p-values correspond to two-way ANOVA of cell density comparing treatment groups and taking zones into account.

We also analyzed these treatment combinations at TP2. PXS-5505 alone did not affect tumor weight, chemo reduced tumor weight but addition of PXS-5505 did not significantly improve this (Figure 6C). However, while chemotherapy alone did not change the number of peritoneal metastatic sites at TP2, there was a significant reduction comparing control and combination therapy (Figure 6D). At end point there was no difference in metastases between the groups (Figure 6E). As with the early treatment, PXS-5505 significantly reduced tumor stiffness, measured by equilibrium and elastic modulus. This was sustained in the combination group (Figure 6F). We stained the TP2 tumors for FN1, VCAN, Col1A1 and M3C. There were no changes in protein density, apart from VCAN which decreased with chemo, a change that was sustained with the combination treatment (Figure S6C). Structural and textural analyses revealed that PXS-5505 treatment led to significant changes in Col1A1, with shorter fibers organized in a less complex pattern (lower BCFD), and an increase in lacunarity, suggestive of increased number and size of gaps in the matrix. Additionally, COL1A1 and FN1 fibers displayed a reduction in curvature, and there was a reduction in HDM in M3C-stained tumors (Figure 6G). Most of the COL1A1 changes were sustained in the combination group as well as the significant changes in FN1 (Figure S6D). Collectively, these alterations of the ECM structure are consistent with the tumor stiffness reduction observed in Figure 6F.

Using multiplex IHC and spatial analysis to study the TME (Figures 6H Figure S6E), we measured the distribution of cells in a 500μm area from the edge to the tumor center. There was a significant increase in CD8+ T cells and B220+ B cells with PXS-5505 treatment at TP2 (Figure 6I). The CD8 increase was also significant in the combination treatment arm compared with chemotherapy alone. TAMs were significantly increased in PXS-5505-treated tumors versus control. In addition, cancer-associated fibroblasts displayed a significant change in their distribution in the combination treatment arm compared to chemotherapy alone, with lower densities observed in the combination treated tumors (Figure 6I).

Therefore, although PXS-5505 alone did not influence mouse survival or metastatic spread, this oral treatment successfully changed the structure and density of collagen, along with other ECM molecules, and reduced tumor stiffness, alterations that were largely retained when PXS-5505 was combined with chemotherapy. In addition, PXS-5505 alone and in combination with chemotherapy increased CD8, B cell and TAM infiltration in the established tumors.

## Discussion

The effects of therapy on the TME are often studied at a single time point, usually after NACT. The results from our study show for the first time the dynamic transcriptional and ECM protein changes over a period of twenty-one days in mouse HGSOC models after three doses of chemotherapy. In 60577, a chemo-sensitive model, chemotherapy altered the structure of several key ECM molecules, resulting in a looser and more open ECM. Chemotherapy induced dynamic changes in ECM regulation, innate and adaptive immune response pathways, at the RNA level. Furthermore, chemotherapy-induced immune pathway changes in 60577 tumors showed significant overlap with those previously reported in responsive human HGSOC patients (19), showing relevance of our mouse experiments to human cancers. In HGS2, a partially responsive model, chemotherapy only induced weak and transient transcriptomic and ECM structure changes. If immune cell signatures were altered during response to chemotherapy, these changes were not maintained.

The anti-TME signature we adapted from Bagaev et al. correlated with omental tumor weight and was therefore a good metric to assess response to chemotherapy in our mouse models. Using this signature, we showed several ECM structure and texture metrics correlated with response to chemotherapy. Some of these also associated with progression free survival in human HGSOC, further supporting our conclusion that ECM structure impacts response to chemotherapy. Combining the metrics associated with chemotherapy response, we computed the structure index, which summarized features of the ECM that correlated with response to chemotherapy. Thus, the structure index was a measurement of chemo-responsive ECM. Matrisome genes whose expression correlated with the structure index were therefore involved in building or maintaining a chemo responsive ECM, and those that were altered in 60577, but not in HGS2, were potential therapeutic targets.

This study reports a list of matrisome gene targets that may be involved in fostering ECM features associated with chemo response. Amongst these, we chose to further investigate LOX and LOXL2, because their mRNA levels associated with poor prognosis in patients and with response to chemotherapy in our mouse models. LOX and LOXL2 were also in an extensive regulatory network of many other ECM-genes in our transcriptional dataset.

Long exposure of HGS2 tumor-bearing mice to a highly selective pan-LOX inhibitor, PXS-5505, from an early stage of development confirmed our hypothesis that preventing build-up of excessive collagen crosslinking would enhance the effect of chemotherapy both in reduction of tumor burden and metastasis at time point and increase survival, although we note this effect was modest. The increase in median survival was consistent with previous studies showing PXS-5505 enhanced the effect of gemcitabine in a mouse model of pancreatic cancer(26), as well as the effect of fluorouracil and oxaliplatin (FOX) in intrahepatic cholangiocarcinoma (30). Unlike the pancreatic cancer study, in our HGS2 model, three doses of carboplatin and paclitaxel treatment did not induce fibrosis. Since PXS-5505 inhibits *de novo* collagen cross-linking, we suggest treatment of established tumors in absence of chemotherapy-induced fibrosis limits the treatment’s efficacy. In addition, PXS-5505 was given *ad libitum* in chow. We observed mice ate less following chemotherapy treatment or at advanced stages of disease progression (data not shown), which could restrict PXS-5505 intake and account for the lack of effect on survival.

However, PXS-5505 had significant effects on the TME that could be relevant to its clinical use. Alone, or given with carboplatin/paclitaxel to established peritoneal tumors, we found significant modifications of ECM structure, concomitant with a decrease in tumor stiffness and an increase in CD8^+^ and B cell infiltrate. This was mitigated by an abundance of fibroblasts and CD163+ macrophages which may have suppressed any anti-tumor activity. When PXS-5505 was given as tumors developed, the distribution and density of TAMs and fibroblasts significantly decreased however, which may have contributed to the improvement in survival with PXS-5505 alone and its enhancement of the chemotherapy response. On the other hand, early PXS-5505 treatment did not increase B and CD8^+^ T cells infiltration. This could be due to an insufficient decrease in elastic modulus, which has previously been associated with immune cell infiltration(28, 29), and we cannot rule out that a transient increase in B and CD8+ T cells occurred at an earlier time point. The reduction in TAMs and fibroblasts in these tumors and the increase in B and T cells in the advanced tumors suggests that this treatment combination has the potential of creating a more favorable tumor microenvironment, where combination with a type of immunotherapy can be of benefit. In support of this, an extensive study in several mouse models showed how another LOX inhibitor improved T cell migration into tumors and combination with anti-PD1 improved survival in a pancreatic cancer model(31). Collectively, the structural and TME changes associated with PXS-5505 suggest a greater penetration of chemotherapy in the tumor, thus an adjuvant PXS-5505 treatment regime could be beneficial to patients. Overall, our results demonstrate that targeting the ECM has the potential to enhance the effect of chemotherapy in HGSOC and that further combination with immunotherapy could be beneficial to patients.

## Acknowledgements

This project was funded by Cancer Research UK program grants A16354 and A25714 (FRB, CB, PK, FL, EM), UKRI Frontier Research grant [number EP/X028704/1] (FRB, BM, JDJ, RCB-R, EM), Wellbeing of Women ELS960, RTF1013, Barts Charity seed fund G-002595, City of London Development Fund CTRQQR-2021\100004 (SE). JH and EM are funded by City of London CRUK Core Award CTRQQR-2021\100004. RM is funded by Barts Charity Grant ECMG1B6R. PXS-5505 was conceived and developed by Pharmaxis. Pharmaxis has been solely responsible for the development of the PXS-5505 compound using their own funds. PXS-5505 was provided free of charge.

We would like to thank the surgeons from Barts Trust, the St. George’s Hospital Trust and the Barts Cancer Institute (BCI) Tissue Bank, Mr. Faisal Karim from The Royal Marsden Hospital, Barts Cancer Institute and Institute of Cancer Research and Mr. Owen Heath from The Royal Marsden Hospital for sample provision. We would like to thank Miss Anna Malliouri, for her preliminary work with TWOMBLI, Dr Luke Gammon and the Blizard Phenotypic Screening Facility for assistance with our work on Cell DIVE, the Biological Services Unit and especially Mr. Jordan Chattenton, the Animal Technician Service at BCI and especially Mr. Colin Pegrum, Mr. James Cormack, Mr. Hagen Schmidt and Miss Harmony Blythin for assistance and support with in vivo experiments. We would also like to thank Nadia Rahman and the BCI Pathology Core, Drs. Linda Hammond and Sam Wallis from the BCI microscopy core facility (CRUK microscopy core service grant at Barts Cancer Institute, Core Award CTRQQR-2021\100004). This research was additionally supported by the BCI Flow Cytometry Core Facility CRUK Flow Cytometry core service grant at Barts Cancer Institute. Most importantly, we express our gratitude to the patients for donating the samples, without which this work would not have been possible.

## Author Contributions

Project design by FL, PK, EM and FRB. EM designed and performed the bioinformatics analyses. FL and PK designed, performed and analyzed the mouse experiments and analyses of mouse and human tissues. EDF and PK performed indentation analysis. FL, PK and JH performed multiplex immunofluorescence on mouse tissues. LP and WJ performed the aorta assay and provided the chow for the mouse experiments with PXS-5505. CB, SE, JDJ, BM and RCBR performed and analyzed *in vitro* experiments. SE, CB, LP, WJ and JFM made intellectual contributions. RM provided patient material and provided intellectual input. FL, PK, EM and FRB wrote the manuscript, and all authors edited and commented on the manuscript.

---

**Figure S1: A.** Hierarchical cluster analysis based on Pearson’s correlation matrix of all genes in 60577 tumors and the complete clustering method. **B**. Barplots illustrating the top significantly enriched Gene Ontology Biological Processes of the genes in the clusters 1 to 7 (broken line indicates adjp = 0.05)

**Figure S2: A.** Matrix patterns for FN1, VCAN, COL1A1 and Masson’s Trichrome (M3C) for 60577 TP1, as they were assessed by TWOMBLI and Haralick feature analysis and visualized in the radar plots. The same selection of TWOMBLI and Haralick metrics as used for TP2 is depicted. dark and light blue show the ECM pattern for control and treated TP1 tumors, respectively. *p<=0.05, **p<= 0.01, ***p<= 0.001, Mann-Whitney test. **B.** Univariate Cox Proportional Hazards Regression Analysis on progression-free survival using TWOMBLI and QuPath image analysis IHC metrics on post NACT samples to split patient groups (human). High and low groups were determined by median values nhigh = 7, nlow = 8. Hazard ratios greater than zero denote increased risk in high group; Hazard ratios lower than zero denote lower risk in high group.

**Figure S3: A.** Hierarchical cluster analysis based on Pearson’s correlation matrix of all genes in HGS2 tumors and the complete clustering method. **B.** Correlation scatter plot of the anti-tumor microenvironment scores with omental tumor weights in 60577. Each dot represents an individual mouse tumor. Points that correspond to different experimental groups are illustrated in different colors. **C.** Venn diagrams illustrating overlap of differentially expressed genes (unadjusted p <= 0.01) in treated versus control at TP1 and TP2 for 60577 and HGS2. **D**. Heatmap of HGS2 unique genes in treated versus control at TP1 and/or TP2. **E.** Representative images of immunohistochemistry for fibronectin (FN1), versican (VCAN), Collagen 1A1 (COL1A1) immunohistochemistry and Masson’s Trichrome (M3C) histochemistry of HGS2 tumors taken at TP2. **F**. ECM patterns for FN1, VCAN, COL1A1 and Masson’s Trichrome (M3C) for HGS2 tumors at TP1, as they were assessed by TWOMBLI and Haralick feature analysis and visualized as radar plots. A selection of TWOMBLI metrics along with Haralick contrast is depicted. The average of structural and textural metrics was calculated and the minimum and maximum range for each metric was computed across all categories. Values were normalized and scaled to the maximum range. Red and orange show the ECM pattern for control and treated TP1 tumors, respectively. **G**. ECM patterns for treated TP1, TP2 and TP3 tumors in Red, salmon and orange, respectively. HDM: High Density Matrix, Gap: Mean Gap Area, BCFD: Box Counting Fractal Dimension, Curvature10 to 40: Curvature metric assessed at windows of 10 to 40 pixels, Branchpoints: normalized branchpoints, Endpoints: normalized endpoints, Fiber Length: Average Fiber length. *p<=0.05, **p<= 0.01, ***p<= 0.001, Mann-Whitney test.

**Figure S4: A.** Stacked barplot illustrating the number of matrisome genes per matrisome category that had a positive or negative association with structure index in 60577 but not in HGS2 (Spearman’s r >= |0.5| and p <= 0.05) **B.** Correlation scatter plots illustrating *Loxl2* correlation with structure index in both mouse models, each dot represents an individual mouse. **C**. Immunohistochemical staining and **D.** Quantification for *LOXL2* in control and chemotherapy treated tumors from 60577 (blue) and HGS2 models (orange). % positive area in the tumor plus stromal area (excluding fat) was quantified in both cases. Each dot represents a mouse, p-values correspond to Kruskal-Wallis test and Mann-Whitney for individual time points * p<0.05, ** p<0.01, *** p<0.001. A representative image from TP2 is shown from each tumor, scale bar: 100um. **E.** Correlation network constructed using the matrisome genes with a positive (pearson’s r>=0.7, p <= 0.05) or negative correlation (pearson’s r>=0.7, p <= 0.05) with LOX. AntiTME signature and structure index circled in fuchsia.

**Figure S5: A**. Quantification of % FN1+, VCAN+, COL1A1+ and M3C+ area for HGS2 biopsies treated with early PXS-5505 in the presence or in the absence of chemotherapy. p-values correspond to two-tailed Mann-Whitney U test. Control n=16-18, Chemo n=16, Early PXS-5505 n=8, Early combo n=9. Each dot represents an individual mouse. **B.** Structural and textural modifications of ECM induced by early treatment with PXS-5505 and chemotherapy in HGS2 tumors at TP2. Radar plots computed as previously, show selected metrics for FN1, VCAN, COL1A1 and M3C fibers. The scaled average for each group is depicted. Control in grey, early PXS-5505 in light blue, Chemo group in yellow. P-values correspond to two-tailed Mann-Whitney U test. Data pooled from two individual experiments are shown, along with controls and chemo treated samples from a third experiment. Control n=16-18, Chemo n=16, Early PXS-5505 n=8. **C.** A representative composite image from each treatment group of HGS2 at TP2 is shown with CD4 (fuschia), FoxP3 (white), endomucin (red), cancer cells (CK14/CK19, green) and COL1A1 (yellow). Hoechst 33342 in blue. E.PXS-5505: early PXS-5505, E.Combo: Early Combo. Scale bar is 100um.

**Supplemental Fig 6. A**. Lysyl oxidase family activity measured in snap frozen aortas determined by fluorometric activity assay. Number of mice were for control *n* = 4, late PXS-5505 n=5, chemo n=4 and late combo n=5. Data presented as mean ± sem. Two-tailed *p* value determined by unpaired, nonparametric *t*-test with a Mann–Whitney *U*-test correction. **B.** Survival curves for the three individual *in vivo* experiments described in Fig 5B and Fig 6B, using early and late treatment with PXS-5505 and six cycles of chemotherapy (carboplatin 20 mg/kg, paclitaxel 10mg/kg, once per week). Median survival for control is 68, 81 and 73.5 days, for the 1^st^, 2^nd^ and 3^rd^ experiment, for chemo 82, 95 and 86.5, for late PXS-5505, 73, 81, and 72, for late combo, 94, 95 and 97, respectively. For early PXS-5505, median survival was 84 and 80 days for the 2^nd^ and 3^rd^ experiment and for early combo 119 and 92 days, respectively. The log rank p value is depicted on the survival curves. The start of the treatment is indicated by the red arrow for chemotherapy, green arrow is for PXS-5505. Number of mice enrolled in each arm is shown in parentheses. **C.** Quantification of % FN1+, VCAN+, COL1A1+ and M3C+ area in HGS2 biopsies treated or not with PXS-5505 and in the presence or in the absence of chemotherapy. p-values correspond to two-tailed Mann-Whitney U test. Control n=16-18, chemo n=16, late PXS-5505 n=8, late combo n=10. Each dot represents an individual mouse. **D.** Structural and textural modifications of ECM induced by late treatment with PXS-5505 in HGS2 tumors at TP2. Radar plots show selected metrics for FN1, VCAN, COL1A1 and M3C fibers. Control is shown in grey, late PXS-5505 in light blue. P-values correspond to two-tailed Mann-Whitney U test. **E.** Representative composite image from each treatment group HGS2 at TP2 is shown with CD4 (fuchsia), FoxP3 (white), endomucin (red), cancer cells (CK14/CK19, green) and COL1A1 (yellow). Hoechst 33342 in blue. Scale bar is 100um.

## List of Supplemental Tables

<u>Table S1.</u> Gene cluster membership, annotation and Gene Ontology Biological Pathways enrichment related to Figure 1.

<u>Table S2.</u> GSVA enrichment scores using the Bagaev et al. PMID: 34019806 functional gene expression signatures, related to Figure 1.

<u>Table S3.</u> Overlapping canonical pathways in 60577 treated vs Control TP1 or TP2 and the Adzibolosu et al. dataset (GEO227100), related to Figure 1.

<u>Table S4.</u> Spearman’s correlation coefficients for ECM genes with Structure Index in 60577 but not in HGS2, related to Figure 4 and S4.

<u>Table S5.</u> Antibodies and conditions for immunohistochemistry, Cell DIVE and ECM TWOMBLI analysis

## List of resources and reagents

| REAGENT or RESOURCE | SOURCE | IDENTIFIER |
| --- | --- | --- |
| Antibodies |  |  |
| Collagen I antibody | Abcam | AB_446161 |
| Rabbit Anti-Collagen I Polyclonal antibody Unconjugated | Abcam | AB_731684 |
| anti-Fibronectin antibody produced in rabbit | Sigma-Aldrich | B_476976 |
| anti-VCAN antibody produced in rabbit | Sigma-Aldrich | B_1080561 |
| anti-LOX antibody (EPR4025) | Abcam | AB_2630343 |
| LOXL2 antibody | Abcam | AB_10677617 |
| Goat Anti-Mouse Mmr Polyclonal antibody, Unconjugated | R&D Systems | AB_2063012 |
| Recombinant Anti-Cytokeratin 14 antibody (EPR17350)-<br>Cytoskeleton Marker | Abcam | AB_2811031 |
| Endomucin (V.7C7) PE conjugated | Santa Cruz | AB_2100037 |
| Col-1A1 | Cell Signalling<br>Technology | Cat# 72827 |
| Recombinant Alexa Fluor 555 Anti-CD8a antibody<br>(EPR21769) | Abcam | AB_2927548 |
| CD4 | Abcam | Cat# ab277275 |
| F4/80 | Cell Signalling<br>Technology | Cat# 25514SF |
| Fibronectin | Abcam | Cat# ab313350 |
| B220 / CD45R | BD Pharmigen | Cat# 553089 |
| Vimentin | Cell Signalling | Cat# 69227 |
| Anti-PCNA antibody (PC10) Alexa Fluor R 647 | Abcam | AB_2857977 |
| FoxP3 | eBioscience | Cat# 12-5773-82 |
| S100A4 | Abcam | Cat# ab208566 |
| aSMA | Novus Biologicals | Cat# NBP2-33006AF750 |
| Perilipin (PLIN1) | Cell Signalling<br>Technology | Cat# 52975S |
| Recombinant Anti-Cytokeratin 19 antibody (EP1580Y)<br>(Alexa Fluor R 555) | Abcam | AB_2857974 |
| CD163 | Abcam | Cat# ab313666 |
| APC anti-human S100A4 | Biolegend | AB_2616691 |
| Rabbit IgG Isotype Control | Thermofisher Scientific | Cat# 31235 |
| Rat IgG2a kappa Isotype Control (eBR2a), eBioscience™ | Thermofisher | Cat# 14-4321-81 |
| Mouse IgG2a kappa Isotype Control (eBM2a),<br>eBioscience™ | Thermofisher | Cat# 14-4724-82 |
| Mouse IgG1 kappa Isotype Control (P3.6.2.8.1),<br>eBioscience™ | Thermofisher | Cat# 14-4714-82 |
| Biotinylated Goat Anti-Rabbit IgG (1ml) | Vector Laboratories | Cat# BA-1000-1.5 |
| Biotinylated Goat Anti-Mouse IgG | Vector Laboratories | Cat# BA-9200-1.5 |
| Horse Anti-Goat IgG Antibody (H+L), Biotinylated | Vector Laboratories | Cat# BA-9500-1.5 |
| Biotinylated Goat Anti-Rat IgG | Vector Laboratories | Cat# BA-9401-.5 |
| Donkey Anti-Goat IgG H&L (Alexa Fluor® 750)<br>preadsorbed. Min X Hu,Mu,Rb,Rat | Abcam | AB_2924800 |
| Donkey anti-Rabbit IgG (H+L) (Alexa Fluor™ Plus 647)<br>Highly Cross-Adsorbed. Min X Gt,Mu,Hu,Rat | Thermo Fisher<br>Scientific | Cat# A32795 |
| Mannose Receptor (CD206) | R&D Systems | Cat# AF2535 |
| Anti-Cytokeratin 14 antibody [EPR17350] - Cytoskeleton<br>Marker | Abcam | Cat# ab181595 |
| Endomucin | Santa Cruz | Cat# sc-65495-PE |
| Col-1A1 | Cell Signalling<br>Technology | Cat# 72827 |
| CD8a | Abcam | Cat# ab280863 |
| CD4 | Abcam | Cat# ab277275 |
| F4/80 | Cell Signalling<br>Technology | Cat# 25514SF |
| Fibronectin | Abcam | Cat# ab313350 |
| B220 / CD45R | BD Pharmigen | Cat# 553089 |
| Vimentin | Cell Signalling | Cat# 69227 |
| PCNA | Abcam | Cat# ab201674 |
| FoxP3 | eBioscience | Cat# 12-5773-82 |
| S100A4 | Abcam | Cat# ab208566 |
| aSMA | Novus Biologicals | Cat# NBP2-33006AF750 |
| Perilipin (PLIN1) | Cell Signalling<br>Technology | Cat# 52975S |
| CK-19 | Abcam | Cat# ab203444 |
| CD163 | Abcam | Cat# ab313666 |
| Rabbit IgG Isotype Control | Thermofisher Scientific | Cat# 31235 |
| Rat IgG2a kappa Isotype Control (eBR2a), eBioscience™ | Thermofisher | Cat# 14-4321-81 |
| Mouse IgG2a kappa Isotype Control (eBM2a),<br>eBioscience™ | Thermofisher | Cat# 14-4724-82 |
| Mouse IgG1 kappa Isotype Control (P3.6.2.8.1),<br>eBioscience™ | Thermofisher | Cat# 14-4714-82 |
| Biotinylated Goat Anti-Rabbit IgG (1ml) | Vector Laboratories | Cat# BA-1000-1.5 |
| Biotinylated Goat Anti-Mouse IgG | Vector Laboratories | Cat# BA-9200-1.5 |
| Horse Anti-Goat IgG Antibody (H+L), Biotinylated | Vector Laboratories | Cat# BA-9500-1.5 |
| Biotinylated Goat Anti-Rat IgG | Vector Laboratories | Cat# BA-9401-.5 |
| Donkey Anti-Goat IgG H&L (Alexa Fluor® 750)<br>preadsorbed. Min X Hu,Mu,Rb,Rat | Abcam | Cat# ab175745 |
| Donkey anti-Rabbit IgG (H+L) (Alexa Fluor™ Plus 647)<br>Highly Cross-Adsorbed. Min X Gt,Mu,Hu,Rat | Thermo Fisher<br>Scientific | Cat# A32795 |
| Bacterial and virus strains |  |  |
| Biological samples |  |  |
| Chemicals, peptides, and recombinant proteins |  |  |
| Xylene, Certified AR | Fisher Scientific Ltd | Cat# X/0250/17 |
| Xylene, Technical | Fisher Chemical Fisher Scientific | Cat# X/0100/17 |
| Ethanol Absolute 99.8+% | Fisher Scientific Ltd | Cat# E/0650/DF/17 |
| Ethanol, 99.8%, as ethanol, anhydrous, (denat .with 2% IPA + 2% MEK) | Fisher Scientific | Cat# 12367103 |
| Antigen Unmasking Solution, Citric Acid Based | Vector Laboratories | Cat# H-3300-250 |
| Antigen Unmasking Solution, Tris Based | Vector Laboratories | Cat# H-3301-250 |
| 10x TBS Buffer pH 7.4 | Severn Biotech Ltd | Cat# 20-7301-10 |
| Hydrogen peroxide solution 30% | Sigma-Aldrich | Cat# H1009-500ml |
| Hydrogen peroxide solution 30% | Fisher Scientific Ltd | Cat# H/1800/15 |
| Antibody diluent | Zytomed Systems | Cat# ZUC025-100 |
| Antibody diluent | Vector Laboratories | Cat# SP-5035 |
| Normal Goat Serum | Sigma-Aldrich | Cat# G9023-10ML |
| Normal Donkey Serum | Merck Life Science Ltd | Cat# S30-100ML |
| Horse Serum (10ml) | Sigma-Aldrich | Cat# H0146-10ML |
| 10X Casein (250ml) | Vector Laboratories | Cat# SP-5020 |
| Sodium Hydroxide 1M | Fluidigm | Cat# J/7620/15 |
| Glycerol | Sigma Aldrich | Cat# G9012-500ML |
| Tween 20 | Sigma Aldrich | Cat# P1379-250ML |
| Hoechst 33342, Trihydrochloride, Trihydrate - FluoroPure™ Grade | Thermofisher | Cat# H21492 |
| Hematoxylin Solution, Gill No. 1 | Sigma -Aldrich | Cat# GHS132 |
| Gill's hematoxylin I | Sigma-Aldrich Cat | Cat# GHS1128 |
| collagenase from Clostridium histolyticum | Sigma-Aldrich | Cat# C9263 |
| DNase I from bovine pancreas | Sigma-Aldrich | Cat# D4513 |
| HBSS (10X), no calcium, no magnesium, no phenol red | GIBCO | Cat# 14185-045 |
| Fixable Viability Dye eFluor 506 | eBioscience | Cat# v65-0866-18 |
| Formalin solution neutral buffered 10% | Sigma Aldrich | Cat# HT501128 |
| Diaminobenzidine substrate-chromogen (Dako Liquid DAB+ Substrate Chromogen System) | Dako | Cat# K3468 |
| DPX Mountant for histology | Sigma-Aldrich | Cat# 6522 |
| Bouin's Solution | Sigma Aldrich | Cat# HT10132 |
| Weigert's Iron Hematoxylin Solution | Sigma Aldrich | Cat# HT1079 |
| Glacial acetic acid | Fisher Scientific | Cat# 10021123 |
| Dubelcco's Phosphate Buffered Saline | Sigma-Aldrich | Cat# D8537 |
| Penicillin-Streptomycin | Sigma | Cat# P4333 |
| L-glutamine | Sigma | Cat# G7513-100ML |
| RPMI1640 | GIBCO | Cat# 21875034 |
| Insulin, Transferrin, Selenium, Sodium Pyruvate Solution (ITS-A) | GIBCO | Cat# 51300044 |
| DMEM:F12 | Gibco | Cat# 31331093 |
| Antigen Retrieval Buffer (100X Tris-EDTA Buffer, pH 9.0) | Abcam | Cat# ab93684 |
| Bovine Serum Albumin | Sigma Aldrich | Cat# A4503 |
| Goat serum | Sigma Aldrich | Cat# G9023 |
| Phosphate-buffered saline (PBS) (1x) | GIBCO | Cat# 141-90-094 |
| Proteinase K in TE-CaCl<sub>2</sub> buffer pH8 | Invitrogen | Cat# 25530015 |
| Tween 20 | Sigma-Aldrich | Cat# P7949-500ml |
| Triton X-100 | Sigma-Aldrich | Cat# T8787 |
| Protein Block, Serum-Free | DAKO Agilent | Cat# X0909 |
| EDTA 0.5M | Thermo fisher | Cat# AM9262 |
| DMEM:F12 1:1 with GlutaMax | GIBCO | Cat# 31331-028 |
| Fetal Bovine Serum | Hyclone | Cat# 11521831 |
| Penicillin-Streptomycin (10,000 U/mL) | Invitrogen | Cat# 15140-122 |
| Hydrocortisone, g-irradiated, powder | Sigma | Cat# H0135 |
| Murine Epidermal Growth Factor (EGF) | Sigma | Cat# E4127 |
| Antibiotic-Antimycotic (100X) | GIBCO | Cat# 5240-062 |
| 0.5% Trypsin-EDTA (10X) | GIBCO | Cat# 15400-054 |
| Carboplatin | Hospira | Cat# 61703-0339 |
| Paclitaxel | Hospira | Cat# 61703-0342 |
| Carboplatin | MedChemExpress | Cat# HY-17393 |
| Paclitaxel | MedChemExpress | Cat# HY-B0015 |
| Fetal Bovine Serum Heat Inactivated | Gibco | Cat# 10438-026 |
| Pan-LOXi | Syntara (previously<br>known as Pharmaxis) | PXS-5505 |
| PXS-5505 mouse diet | Research Diets | Cat# D21012504R |
| Control mouse diet | Research Diets | Cat# D20011301 |
| Koliphor EL | Merck Life Science Ltd | Cat# C5135 |
| Distilled Water | Gibco | Cat# 15230204 |
| Ethanol denatured (BioUltra) | Sigma-Aldrich | Cat# 51976-500ML-F |
| RNAlater-ice | Thermo Fisher<br>Scientific | Cat# 4427575 |
| Critical commercial assays |  |  |
| Avidin/Biotin Blocking Kit | Vector Laboratories | Cat# SP-2001 |
| FlexAble CoraLite® Plus 750 Antibody Labeling Kit for Rabbit IgG | ProteinTech/FlexAble | Cat# KFA004 |
| DAB Peroxidase (HRP) Substrate Kit | Vector Labs | Cat# SK-4100 |
| VECTASTAIN® ABC HRP Kit | Vector Labs | Cat# PK-4000 |
| Trichrome Stain (Masson) Kit | Sigma-Aldrich | Cat# HT15-1KT |
| anti-rat ImmPRESS-HRP reagent | Vector Laboratories | Cat# MP7444 |
| anti-rabbit ImmPress HRP | Vector Laboratories | Cat# MP7451 |
| Super Sensitive Polymer HRP IHC | BioGenex | Cat# QD430-XAKE |
| RNeasy Mini kit | Qiagen | Cat# 74104 |
| QIAshredder | Qiagen | Cat# 79654 |
| High Sensitivity RNA ScreenTape | Agilent | Cat# 5067-5579 |
| MycoAlert PLUS Mycoplasma Detection kit | Lonza | Cat# LT07-710 |
| Deposited data |  |  |
| Chemotherapy-induced extracellular matrix remodelling in HGSOC | GEO | GSE270878 |
| Experimental models: Cell lines |  |  |
| 60577 | Laboratory of<br>Difilippantonio S. | Szabova et al., 2014 |
| HGS2 | Laboratory of Frances Balkwill | Maniati et al., 2020 |
| Experimental models: Organisms/strains |  |  |
| FVB mice | Charles River |  |
| C57/Bl6J mice | Charles River |  |
| Oligonucleotides |  |  |
| Recombinant DNA |  |  |
| Software and algorithms |  |  |
| HALO | Indica labs |  |
| FIJI | PMID: 22743772 | https://fiji.sc/ |
| QuPath version 0.2.0 to 0.5.0 | https://qupath.github.io/ | Bankhead et al., 2017) |
| FlowJo software, Tree Star, Inc. FlowJo Software for PC | BD Bioscience | v.10.1.5 |
| Version 10.Ashland |  |  |
| NPD.view 2.7.25 software<br>[https://www.hamamatsu.com/eu/en/](https://www.hamamatsu.com/eu/en/product/type/U12388-01/index.html) |  |  |
| [product/type/U12388-01/index.html](https://www.hamamatsu.com/eu/en/product/type/U12388-01/index.html) |  |  |
| ImageJ 1.48v, 1.54f | NIH, Bethesda, MD,<br>Schneider et al., 2012;<br>Ruifrok and Johnston, 2001 | https://imagej.net/ |
| TWOMBLI plugin |  |  |
| imagej.net/Colour_Deconvolution |  |  |
| Definiens software | (Definiens AG,<br>Germany) | http://www.astrazeneca.com |
| Graphpad Prism, San Diego, CA | NA | https://www.graphpad.com |
| R v4.1 | NA | http://www.R-project.org |
| EdgeR | Bioconductor | Robinson MD et al., 2010 |
| limma | Bioconductor | Ritchie ME et al., 2015 |
| GSVA | Bioconductor | Hänzelmann S et al., 2013 |
| biomaRt | Bioconductor | Durinck S et al., 2009 |
| fgsea | Bioconductor | Korotkevich G et al., 2019 |
| ComplexHeatmap | Bioconductor | Gu Z et al., 2016 |
| ggplot2 | CRAN | https://cran.r-project.org/web/packages/ggplot2/index.html |
| gplots | CRAN | https://cran.r-project.org/web/packages/gplots/index.html |
| Matlab | MathWorks | https://www.mathworks.com/products/matlab.html |
| Other |  |  |

## Methods

### Experimental model and study participant details

#### Study approval

##### Murine models

All *in vivo* experimental procedures observed the guidelines approved by the ethics committees of QMUL in accordance with Animals (Scientific Procedures) Act 1986 under the Home Office Project licenses PBE3719B3 and PP5394401. For survival experiments, mice were euthanized when they reached humane endpoint as defined in the licenses.

##### Patient samples

Samples were kindly donated by HGSOC patients undergoing surgery at Barts Health NHS Trust, the Royal Marsden Hospital and St. George’s University Hospitals NHS Foundation Trust. Tissues deemed by a pathologist to be surplus to diagnostic and therapeutic requirements were collected along with clinical data under the Barts Gynae Tissue Bank HTA license number 12199 (REC no: 10/H0304/14, 15/EE/0151, from the Royal London Hospital (ethics number: 17/LO/0405) and from the Royal Marsden Hospital (REC no: 20/LO/0529). Patients gave written informed consent, and the study was approved by a UK national review board. Studies were conducted in accordance with the Declaration of Helsinki and the International Ethical Guidelines for Biomedical Research Involving Human Subjects.

#### Animals, *in vivo* models, treatment, and tissue processing

For the orthotopic tumor growth establishment, 60577 and HGS2 mouse cell lines were trypsinized, washed in medium and resuspended in PBS to 10^7^ cells in 300μl to be injected intraperitoneally (i.p.) in 8-9-week-old FVB mice (60577) or C57BL/6J mice both from Charles River (UK). For chemotherapy treatment, mice were treated with a combination of carboplatin (20 mg/kg, Hospira) + paclitaxel (10 mg/kg, Hospira), both from the pharmacy at St. Bartholomew’s Hospital, London or from Medchemexpress (HY-17393 and HY-B0015, respectively). Carboplatin and paclitaxel were administered to mice via i.p. in 200 μL volume starting respectively 21 days (60577), or 49 days (HGS2) post-cell injection. Vehicle-treated controls received Koliphor® EL (Merck, Cat# C5135), 6.1%, EtOH 6.1%, water 14.7%, PBS 73.1%. Mice were treated with chemotherapy or vehicle via i.p. once weekly for 3 (TP2) to 6 weeks.

For Pan-LOX inhibition (LOXi) in the HGS2 model, we used PXS-5505 (Syntara) under an MTA agreement. PXS-5505 was provided in the chow (D21012504R, Research Diets) ad libitum in a concentration that is approximately 170mg/kg (maximum dose calculated). The control group mice, control chow D20011301 (Research Diets) was provided. After one week of acclimatisation, all mice were introduced to Control chow, D20011301. Mice were then introduced to D21012504R PXS-5505 diet after 7 and 49 days for the early and late PXS-5505 treatment, respectively. PXS-5505 treatment continued up to 22 weeks. For combinatorial treatment of PXS-5505 with chemotherapy in HGS2 model, chemotherapy treatment commenced described as above.

Humane endpoint for mice was defined as a change in general health; specifically, 15% body weight loss over 72 hours or 20% over any time period, any sign of jaundice, hunched posture, or abnormal breathing pattern, signs of distress and/or suffering, as well as signs of ascites or palpable tumors exceeding the license limits. Any tumor interfering with mobility and access to food and water, as well as ulceration or increased secretion of injection site tumors, blood in the stool and or urine defined humane endpoint additionally. Survival assessment of mice was made by the same individual where possible to limit inter-observer variability and occasionally assistance was provided by a trained animal technician who was not directly involved in the experimental design.

For both time point and endpoint analysis, mice were euthanized under CO2 exposure. Once death was confirmed, an incision to expose the gastrointestinal contents was made that was extended further towards the lower limbs bilaterally to create skin flaps which can be pinned down. Omental, splenoportal and lesser omental tumors were harvested, along with mesentery and their weights measured. Omental tumors were cut in half, with one half being snap frozen in dry ice and the other half fixed in 10% neutral buffered Formalin solution overnight. The formalin fixed tumors were paraffin-embedded and sectioned (4 μm), followed by H&E staining, whereas the frozen tumors were subjected to RNAseq or indentation testing. Tumors in other anatomical sites were excised and weighed. Once all tumors were recorded and weighed, internal organs like liver, pancreas, stomach, spleen, and intestines were removed to better visualize the aorta that was subsequently separated from the spine dorsally and the oesophagus ventrally. Once isolated, aortas were snap frozen in dry ice and kept at −80C.

#### Murine cell line culture and *in vitro* chemoresistance

HGS2 and 60577 described previously(13) were cultured in complete DMEM:F12 1:1 with GlutaMax medium (GIBCO) with the addition of 4% heat inactivated FBS (Hyclone, Gibco), 1x Pen/Strep (Invitrogen), 1x insulin/transferrin/selenium (Invitrogen, 51300), 100 ng/ml hydrocortisone (Sigma, H0135), 20 ng/ml murine Epidermal Growth Factor (EGF, Sigma, E4127), 1x antibiotic-antimycotic (GIBCO). Cells were trypsinized with 0.05% trypsin-EDTA (GIBCO) and regularly tested for Mycoplasma, using the MycoAlert PLUS Mycoplasma Detection Kit (cat. no. LT07-710, Lonza). For the chemosensitivity assays, the day prior to stimulation, 60577 and HGS2 cells were plated at a density of 20,000 cells/well, in a 24 well plate, in triplicates. The day after, cells were stimulated with 1, 0.5, 0.1, 0.01, or 0.001mM of carboplatin (Hospira, Cat# 61703-0339), and 1, 0.5, 0.1, 0.01 or 0.001uM paclitaxel (Hospira, Cat# 61703-0342). At 48h, cells were harvested with trypsin and counted with a V-cell-counter (VI-CELL XR Cell Viability Analyzer: BMSB J540). The percentage live cells compared to control was calculated.

#### Lentivirus transduction of the murine cell lines to induce stable expression of nuclear red protein, mKate

IncuCyte NucLight Red Lentivirus reagent (Essence bioscience, Cat no.: 4475) was used to induce a stable expression of the red mKate2 protein in the cancer cells. The recommended multiplicity of infection (MOI) ranges between 3 to 6. The viral titre was 11.47x 106 transduction units (TU) /ml (Lot number: LSP032219.02-051019). 10,000 cells were plated in a 48 well plate and left to grow until they became 30% confluent. The transduction was started by replacing the conditioned medium in the well with 1 mL of transduction medium that contained 3.5μL of lentiviral particles. Cells were cultured for 24 h in an incubator at 37 °C, 95% humidity and 5% CO2. After 24h, the transduction medium was replaced with normal medium and the cells were cultured for 48-72h in an incubator at 37 °C, 95% humidity, and 5% CO2. The lentivirus integrates into the DNA of the cell and produces mKate2 protein which is red fluorescent. Transduced stable cell populations expressing mkate2 protein were generated by puromycin selection. To determine the dose of puromycin, 250,000 cells were plated per well in 12 well plate for 9 wells. One well was used as control well and the cells in the other 8 wells were treated with increasing concentrations of puromycin starting with 0.5μg/ml to 4μg/ml daily for 4 days. The lowest puromycin concentration that killed all cells in the well was chosen for puromycin selection. For human cell lines 2μg/ml was used. A dose of 1.5μg/ml was chosen for HGS2 and 4μg/ml for 60577.Lentivirus transduced cell lines were stored in 1ml of (10% DMSO+90%FBS) incryovial at −80C then transferred to liquid nitrogen.

#### Incucyte S3 to study the proliferation of murine cell lines

Red labelled lentivirus transduced 60577 and HGS2 were plated as 20,000 cells per well in 2ml of medium in 12 well plate in duplicates. Cells are left to attach for 3-4h then the plate is moved to be imaged in Incucyte S3 for 72h. 4 images per well were taken every 2h. Red object count per well normalized to the counts at 0h were analyzed using incucyte S3 software. Data was then exported from the software and used in Prism version 10.10) to generate AUC for each individual well. The AUC was then used to study statistical significance using unpaired t test.

#### Lysyl oxidase family activity from the mouse aorta

Lysyl oxidases family activity was measured in the tumor bearing aortas of mice at sacrifice. The aorta underwent an urea based extraction(32) and then the level of lysyl oxidase activity was measured with a fluorometric activity assay(33) to confirm lysyl oxidase inhibition. The signal was defined as the enzyme activity in the aorta, while noise is measured in the presence of a high concentration (300 µM) of BAPN to abolish any lysyl oxidase family activity. Signal over noise equal to 1 indicates complete inhibition.

#### Metastasis Analysis

Both at timepoint (TP2) and endpoint, mice were assessed for the number of the metastatic sites identified during autopsy. Main metastatic sites recorded included splenoportal fat, lesser omentum, mesentery, peritoneum, perirenal, spinal, liver, diaphragm, ovaries, ovarian fat pad, fallopian tubes, uterus, uterine fat pad, abdominal fat, chest cavity, gastrointestinal tract. Macroscopical pancreas invasion was also counted as a metastatic site. Each metastatic site counts as 1, regardless of the numbers and size of the metastases on that site. The number of total metastatic sites per mouse was calculated and plotted.

#### Histopathology, immunohistochemistry, and morphometry

Omental tumors from both models were dissected, fixed in 4% formaldehyde for 24 hr, paraffin-embedded and sectioned (4 μm), followed by H&E staining. For immunohistochemistry, 4 μm FFPE omental sections were heated for 1hr at 60°C and then submerged twice in xylene for 5min. Slides were then gradually re-hydrated by submerging for 2min in each of the following ethanol solutions: 100%, 90%, 70%, 50% and finally in ddH2O for 3min. Antigen retrieval was performed as per Table S5A. for all staining apart from FN1. The slides were subsequently washed, treated with 3% H2O2 (Fisher Scientific, H/1800/15) in PBS for 5min, washed again and blocked with blocking buffer for 1hr. The primary antibody was added in blocking buffer and incubated at ambient temperature (Table S5A). Slides were washed three times, and the primary antibody was detected as described in (Table S5A). Color was developed with Diaminobenzidine substrate-chromogen (Dako Liquid DAB+ Substrate Chromogen System, K3468 Dako) and tissues were counterstained with Gill’s hematoxylin I (Sigma-Aldrich, GHS1128), washed, dehydrated in ethanol and xylene, and mounted in DPX mountant (Sigma-Aldrich, 06522). For COL1A1 staining on mouse tissues, ab21286 from Abcam was used, whereas ab34710 (Abcam) was used for staining COL1A1 on human tissues.

#### Masson’s Trichrome staining

To reveal fibrillar collagen deposition on mouse and human omental biopsies, we used the Trichrome Stain (Masson) Kit (HT15-1KT) from Sigma as previously described in(^13^). Briefly, FFPE 4-μm sections were deparaffinized in xylene and gradually re-hydrated by submerging in a graded series of Ethanol solutions in H2O. Slides were further hydrated for 3min in ddH2O and fixed in Bouin’s solution overnight at RT. The next day, sections were washed in running water and ddH2O, until the yellow colour disappeared and counterstained in working Weigert’s Iron Hematoxylin Solution (Sigma, HT1079-1SET). Subsequently, sections were washed in running tap water for 2 minutes, rinsed in ddH2O and stained in Biebrich Scarlet-Acid Fucshin for 15 minutes. Sections were then rinsed in ddH2O and immersed in Working Phosphotungstic/Phosphomolybdic Acid solution for 10-15 minutes, until collagen fibers were not red. Aniline Blue solution was applied next for 30 minutes; sections were rinsed in ddH2O and placed in 1% acetic acid for 3 minutes. Finally, sections were dehydrated very quickly in two changes of 90% alcohol, followed by 2 changes of absolute alcohol, cleared in xylene and mounted in DPX (Sigma-Aldrich, 06522).

#### *In vitro* PXS-5505 treatment experiments

Primary human omental fibroblasts were isolated from macroscopically normal omentum from patients undergoing gynecological surgery, following the protocol previously described(34). Primary human fibroblast were cultured in DMEM:F12 (Gibco, Cat. 31331093) supplemented with 10% FBS, 1% penicillin/streptomycin (Sigma, Cat. P4333), 1% L-glutamine (Sigma, G7513-100ML). For the 2D and 3D experiments, all the cells and gels were maintained in DMEM:F12 with 1% L-Glutamine, 1% ITS (Gibco, Cat. 51300044), 1% penicillin/streptomycin and 4% human serum (Sigma, Cat. H4522). The human malignant HGSOC cell line was labelled with mCherry lentivirus (BPS-Bioscience #78932-P), according to the manufacturers’ instructions. Collagen gels were setup as previously described(34) using 100,000 malignant cells and 100,000 primary fibroblasts. 2D cell lines and 3D collagen gels were treated at day 1 and day 4 with 10 µM PXS-5505.

#### RNA isolation and sequencing

Total RNA was extracted from frozen murine omental tumor samples. The samples were transferred into RNAlater and homogenized with Miltenyi GentleMACS in RLT buffer and further processed using QIAGEN Rneasy Mini kit with on-column Dnase digestion. RNA quality was analyzed with the 4200 TapeStation System using the Agilent High Sensitivity ScreenTape Assay. Library preparation and RNASeq were performed by the Wellcome Trust Centre (Oxford, UK) using NEBNext rRNA Depletion Kit v2 to deplete rRNA species.

Sequencing was performed to ∼65x mean depth on the Illumina NovaSeq6000 platform, strand-specific, generating 150 bp paired-end reads. RNA-Seq reads were quality trimmed using trimgalore v0.6.5 and were mapped to the mouse genome (mm10, Genome Reference Consorstium GRCm38) using STAR v2.7.0f(35). Number of reads aligned to the reference genome were counted using rsem v1.3.1 based on the Ensembl annotation GRCm38 (36). Only genes that achieved at least one read count per million reads (cpm) in at least twenty-five percent of the samples were kept. This led to 16,400 filtered genes in total, 14,031 of which were protein coding genes. Conditional quantile normalization was performed accounting for gene length and GC content and a log2-transformed RPKM expression matrix was generated.

#### Bioinformatic analysis

Differential expression analysis was performed in Edge R using limma(37). Gene set enrichment analysis was performed for Canonical Pathways (c2.cp.v7.4.symbols.gmt) using R package fgsea and the ranked t-statistic of all sufficiently detected orthologous genes. Enrichment analysis of gene clusters or overlapping genes of interest (over-representation analysis) was performed with R package dnet, using a hypergeometric test. Single sample gene-set enrichment analysis which calculates a gene-set enrichment score per sample was performed using the R package GSVA(38). Heatmaps were plotted with R package ComplexHeatmap; scatter plots, barplots and boxplots with R packages gplots and ggplot2. Structure index was obtained by scaling the metrics and calculating the ratio of those parameters that positively correlated with the anti-tumor microenvironment enrichment score over those that negatively correlated with it. The GSE227100 dataset containing RNASeq data of 24 patients with paired pre-NACT and post-NACT RNASeq was downloaded from GEO. All patients had initial tumor biopsies prior to beginning chemotherapy with six cycles of carboplatin and paclitaxel. The patients subsequently underwent post-chemotherapy surgery, which provided the post-NACT samples. All the samples were obtained from metastatic sites, late recurrence samples n = 11. GSEA on this dataset was performed using the ranked log2 fold-change of late post vs pre chemotherapy samples and Canonical Pathways (c2.cp.v7.4.symbols.gmt).

#### Indentation testing of mouse omental tumors

Mechanical characterization of the specimens was conducted employing an Instron 3342 screwdriver mechanical testing frame (Instron, UK) with a crosshead position resolution of 1 μm. The testing apparatus was equipped with a 10 N load cell (resolution of 0.1 mN) and a flat-ended stainless steel cylindrical probe with a 1 mm diameter for conducting indentation tests. Prior to experimentation, the tissue samples were completely thawed at room temperature. During the entire testing duration, the specimens were fully immersed in phosphate-buffered saline (PBS) at room temperature to maintain full hydration. A pre-load of 1mN was applied to the samples, followed by a deformation to 20% at a constant rate of 1%/s. Subsequently, a displacement-hold period of 600 s was implemented. Data analysis was carried out in accordance with the methodology outlined by Delaine-Smith et al. (39), utilizing the mathematical model M2 proposed by Oliver and Pharr (1992), assuming isotropy in the samples and negligible friction. Throughout the analytical process, a set of well-established parameters were computed, including tangent moduli (TM) within the ranges of 2.5–7.5% and 15–20%, as well as peak modulus (PM), equilibrium modulus (EQM), and percentage of Relaxation. Raw data was processed using Matlab and plotted in GraphPad Prism.

#### Multiplex Fluorescent immunohistochemistry (MxIF) staining on mouse HGS2 tumors

For the multiplex immunohistochemistry staining of mouse omental tumors, we used the tissue-cyclic fluorescence (T-CyCIF) methodology(40) adapted to the Cell DIVE™ (Leica Microsystems) imaging platform(41). For the MxIF, serial rounds of conventional immunofluorescence (IF) were facilitated using a fluorophore inactivation step to capture multiple up to 5-plex images on single tissue sections per round for automated downstream registration and assembly into a single high-plex image stack, [PMID: 23818604). 5µm thick FFPE tissue sections were transferred onto SuperFrost Plus™ Adhesion slides (Fisher Scientific) and baked overnight at 60°C. Tissues then were deparaffinized in xylene and rehydrated in graded washes in ethanol to ddH20. Heat-induced epitope retrieval (HIER) was standardized to a pressure cooker cycle of 116°C for 12 minutes with 1X tris-based antigen unmasking solution (H-3301, Vector) for all antibodies used in the panel. Slides were subsequently incubated for 1h at RT in blocking buffer (10% goat serum/1X Casein/TBST 0.1%), followed by a 60 min incubation at RT in a highly oxidative solution (4.5% H2O2 and 20mM NaOH in TBS) in the presence of white light. Two LED light sources, emitting 32,000 LUX each, placed on top and on the bottom of the slide tray were used for this photo-bleach / dye Inactivation procedure that aids in tissue auto fluorescence elimination. Upon completion of the bleaching step, the slides were counter stained with Hoeschst 33342 (5 µg/mL in TBS) (H21492, Thermofisher) for 10 mins before registering the background” images on the Cell DIVE scanner. After image acquisition, slides were washed and incubated overnight at 4C with the antibody cocktail of the first round. Primary antibodies were either validated by the supplier or against both positive mouse tissue controls and disease-related orthotopic HGS2 mouse model biopsies, using isotype (Table S5C) and secondary-only negative controls in place to gage and subtract non-specific staining. All antibodies used in this study were direct conjugates, apart from F4/80, CD206 and KRT14. F4/80 was conjugated to CoraLite® Plus 750, using the FlexAble CoraLite® Plus 750 Labelling kit (KFA004) following manufacturers’ instructions. Indirect detection was achieved via tissue incubation (30 min, R.T) in primary Ab species-specific secondary Donkey IgG’s conjugated to either AF647 or AF750 dye. Fluorescently tagged secondaries (2’Ab) donkey anti-goat IgG-AF750 (H&L) (ab175745, Abcam) and donkey anti-rabbit IgG-AF647 (A32795, Thermofisher) were used in conjunction with purified primary antibodies anti-MMR/CD206 Polyclonal (AF2535, R&D Systems) and anti-Cytokeratin 14 clone EPR17350 (ab181595, Abcam), respectively. The full antibody list for the 17-marker panel we used, along with information on dilution used and staining round composition is shown in detail in Table S5D. The next day, slides were once again incubated with Hoechst33342 staining for 5 minutes, washed and then mounted onto ClickWell Assembly cassettes (Leica Microsystems) in 50% glycerol:TBS (vol:vol) imaging solution. Slides were imaged on the CellDIVE platform, using up to four channels plus counterstain with auto fluorescence removal, corrections and stitching, after which the dye inactivation/antibody staining cycling process was reinitiated. For staining rounds 2-6 bleaching was for 45min. Imaging rounds were conducted over a period of 9 days with an intermediate pause of four months for the late PXS-5505 treatment experiment and 13 days for the early PXS-5505 treatment experiment. Slides were stored for long term at 4C for future experiments.

## Quantification and statistical analysis

### Immunohistochemistry quantification, Haralick feature computation

For standard immunohistochemistry and H&E. tissue sections were scanned with Hamamatsu NanoZoomer S210 Slide-Scanner and the scans analyzed using Definiens® software (Definiens AG, Germany) and QuPath(42). QuPath was used to quantify, FN1, VCAN, COL1A1, M3C and LOXL2 immunohistochemistry staining. Difference of Gaussian (DoG) superpixel classification was used to discriminate between adipose and non-adipose tissue. All staining presented were quantified with positive area thresholders. For FN1, VCAN, COL1A1 and M3C staining, a detection area was created out of the positive pixels detected by the thresholder. The Haralick features were computed in the detection area. Results displayed are measurements of non-adipose tissue. As IHC staining of the different mouse experiments occurred on different days, FN1 and M3C staining displayed noticeable batch effect, with the intensity of one or two experiments out of three clearly different from the others. We therefore normalized the Haralick features (computed based on pixel intensity) using the ratio of the medians of the control groups, using the first experiment as the reference.

### ECM structure quantification using TWOMBLI

ECM structure was analyzed using TWOMBLI(20) a plugin for FIJI (www.imagej.net). Representative images of WSI sections that were stained with Masson’s trichrome, COL1A1, FN1 and VCAN were extracted under x20 magnification as .tiffs and underwent a color deconvolution step before analysis by TWOMBLI, using the Color deconvolution plugin in FIJI. For Masson’s Trichrome, color deconvolution using the H PAS vector in FIJI, allowed for the isolation of blue collagen from the image, whereas for the remaining staining, H DAB vector was used to isolate the brown staining that corresponded to Col1A1, FN1 or VCAN. The parameters used for all different staining for both mouse and human samples are shown in Supplement Table 2. Branch points and endpoints were normalized by dividing by total length; the average fiber thickness was calculated by dividing the high-density matrix by the total fiber length. In Table S5B the exact parameters for each staining are shown, both for mouse and human. Branch points and endpoints were normalized by dividing by total length; the average fiber thickness was calculated by multiplying high-density matrix with image area and dividing by the total fiber length.

### Multiplex Immunofluorescence image acquisition and quantification

Up to 5-plex immunofluorescent Images were acquired with the CELLDIVE ™ (Leica Microsystems) MxIF platform at 20X. Fully stitched Cell Dive images were imported, fused, segmented and analyzed using the HALO® (indica Labs) software. Tissues were classified to segregate adipose, immune aggregates and “tumor” area (non-adipose, excluding immune aggregates). These classifications were masked as annotations and within each along with within the whole tissue, all markers were quantified with HighPlex FL and Area Quantification FL algorithms. This allowed the identification of phenotypes: cytotoxic T cells (CD8+), CD4+ cells, B cells (B220+), FoxP3+ cells, TAMs (CD206+ and/or F4/80+ and/or CD163+ and aSMA-), CAFs (aSMA+ and or S100A4+ and CK14- and CK19- and F4/80-). In addition, to quantify the density and distribution of the different cell types regarding the Tumor area, Infiltration Analysis was run for all phenotypes with the whole tissue as Tissue Annotation Layer and using the Tumor annotation as the Interface layer, with a 500um range and 25 bands.

### Statistical Analysis

GraphPad Prim 10 and R v4.1 were used for statistical analyses. Two-sided non-parametric t-test (Mann-Whitney) was used to assess differences between conditions. When more than two groups were compared, we used non-parametric one-way ANOVA (Kruskall-Wallis). For the assessment of the effect of two factors (for the spatial analysis) we used a two-way ANOVA. Statistical tests used, n numbers and p values are displayed in the appropriate figures and figure legends. P values < 0.05 were considered statistically significant

