## Supplementary figures and images for "Matrix structure and microenvironment dynamics correlate with chemotherapy response in ovarian cancer"

### Supplemental Figure 1

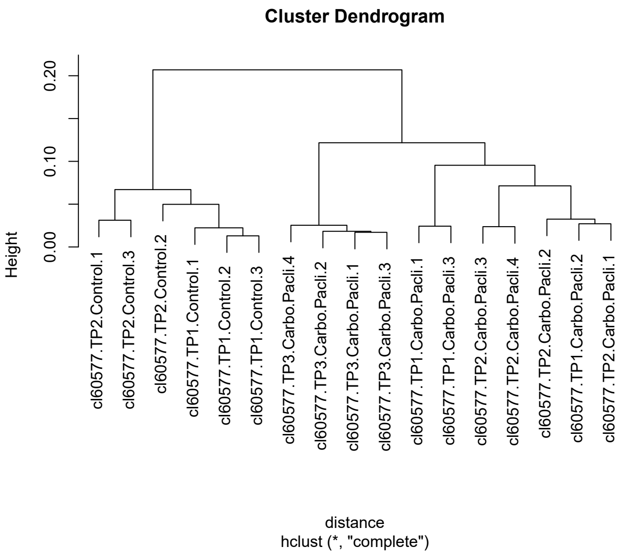

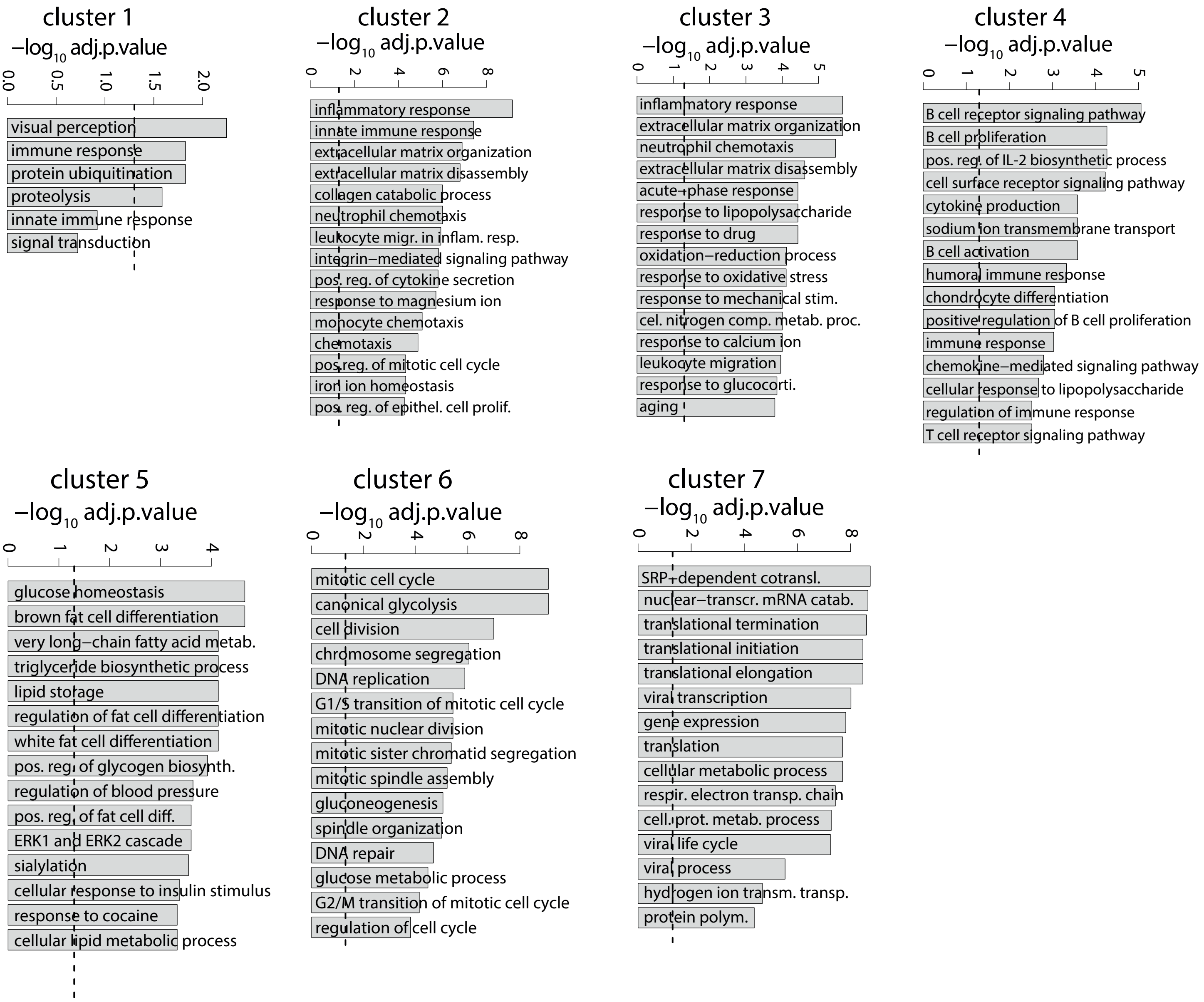

### Supplemental Figure 2

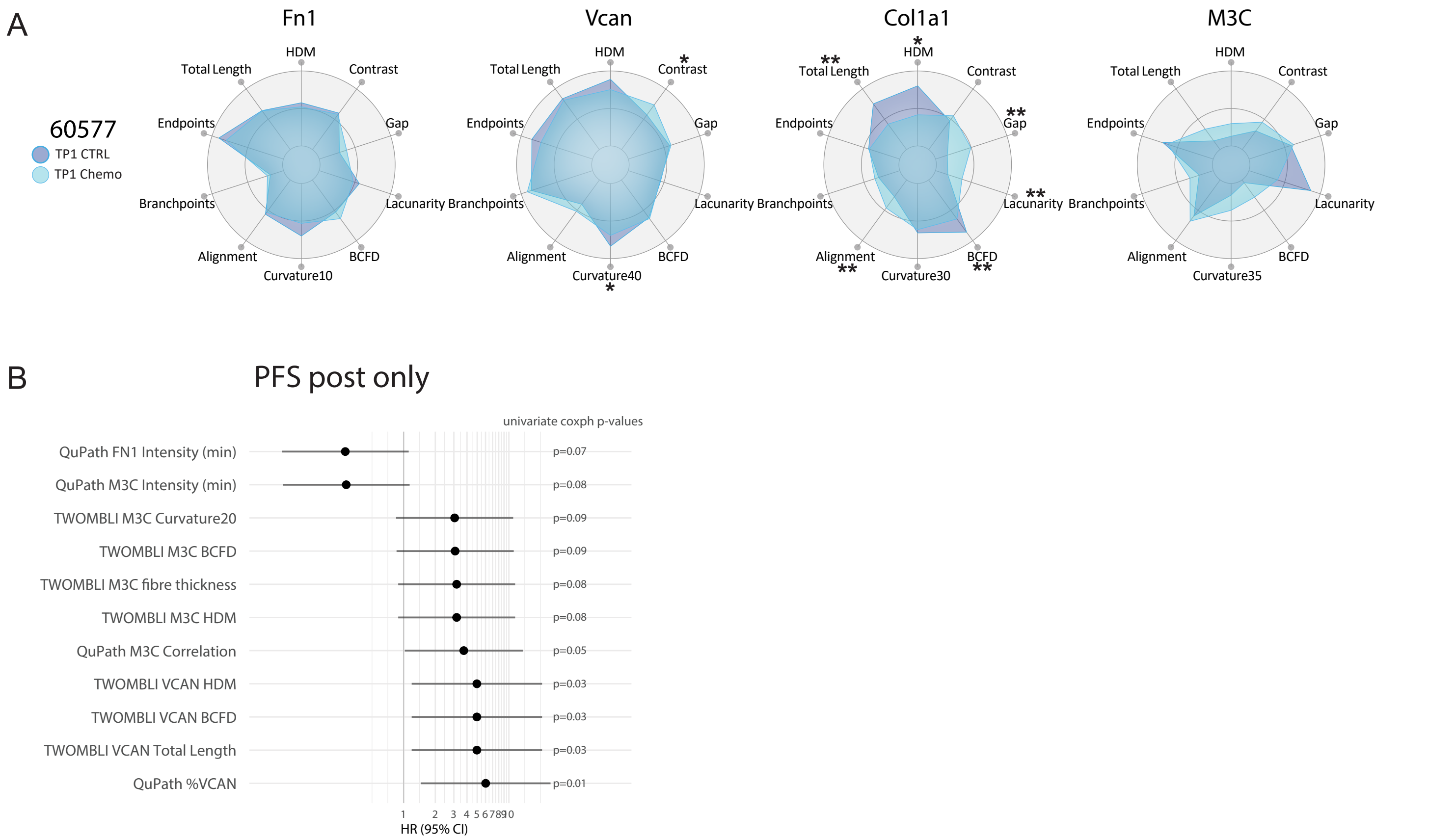

### Supplemental Figure 3

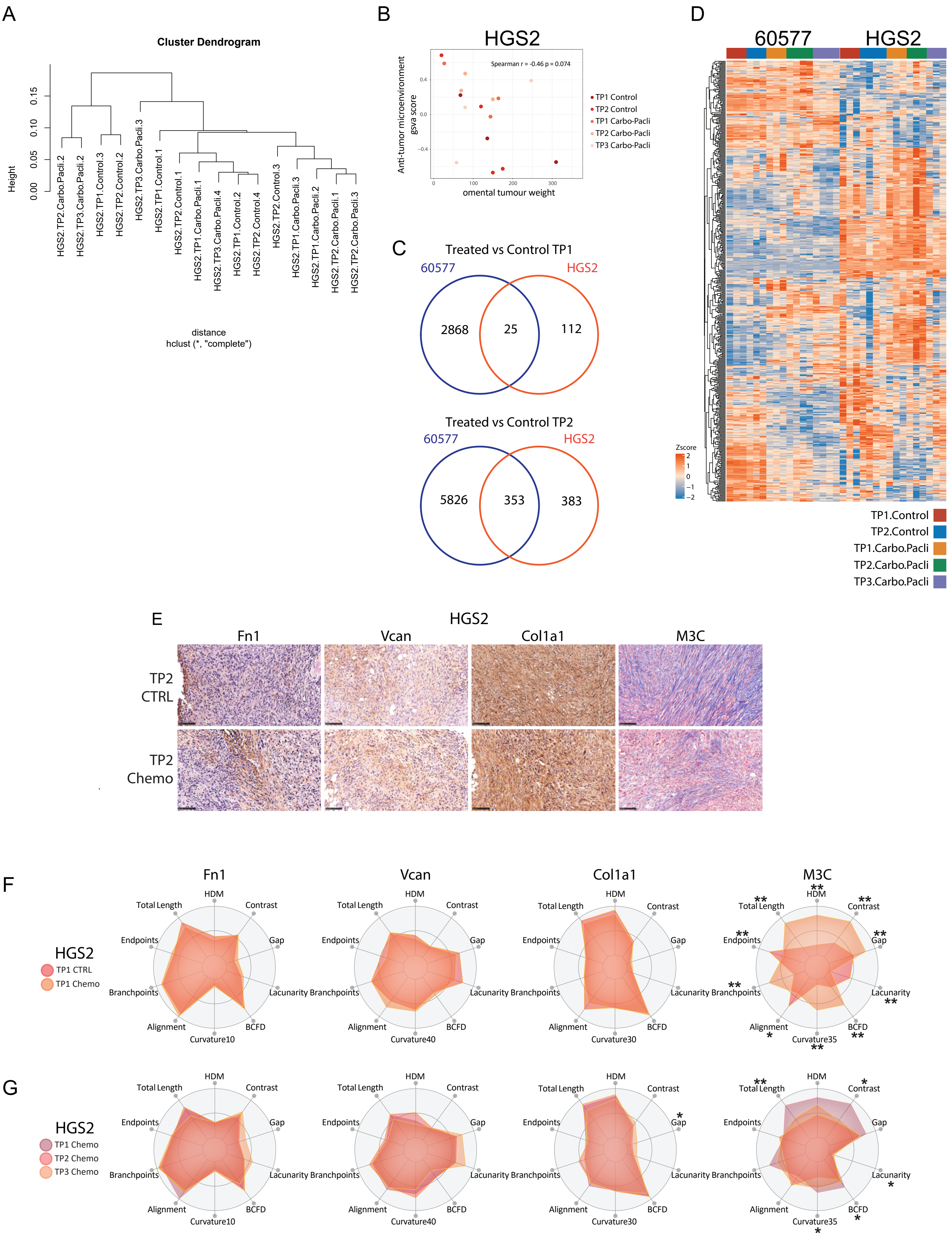

### Supplemental Figure 4

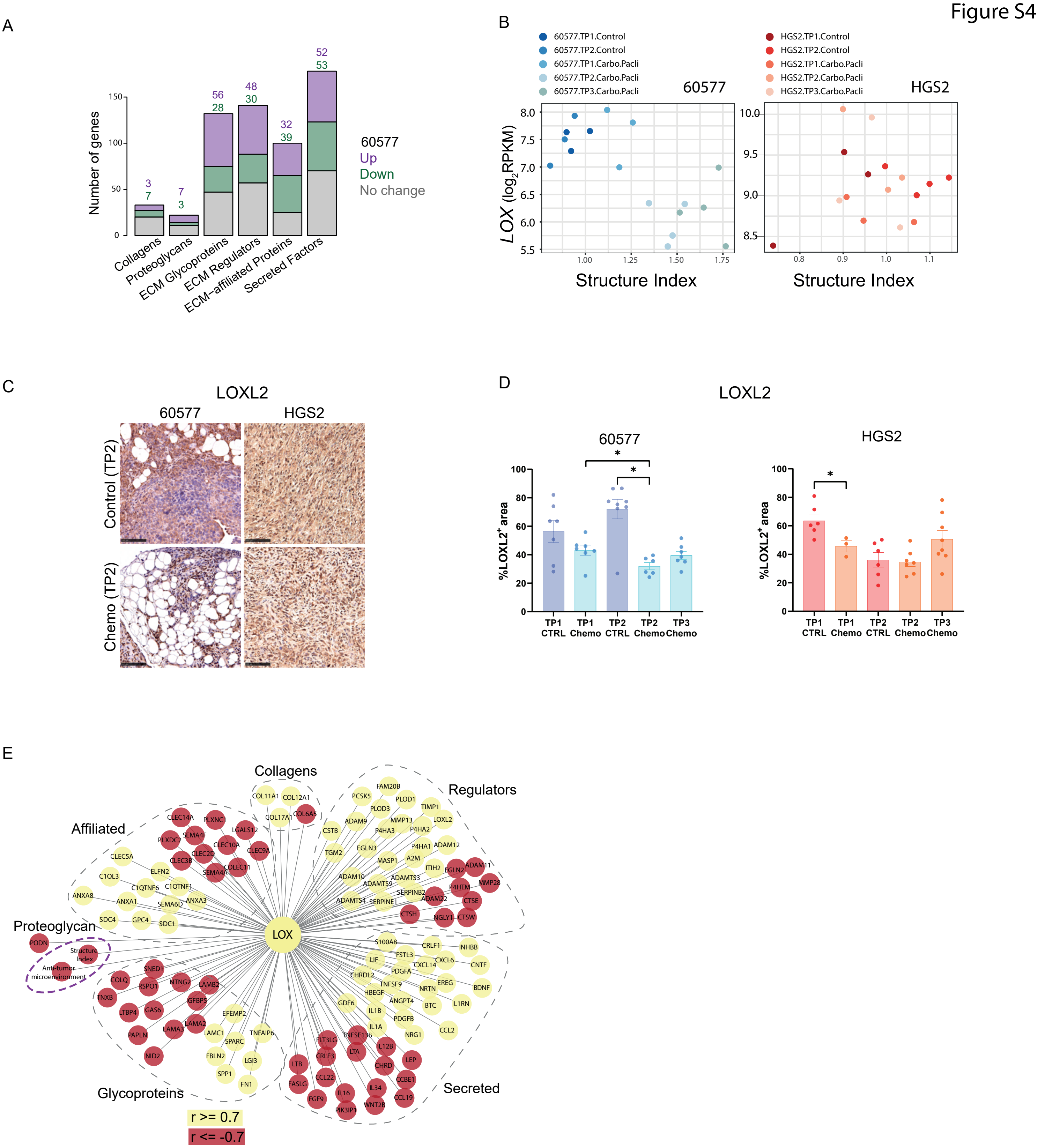

### Supplemental Figure 5

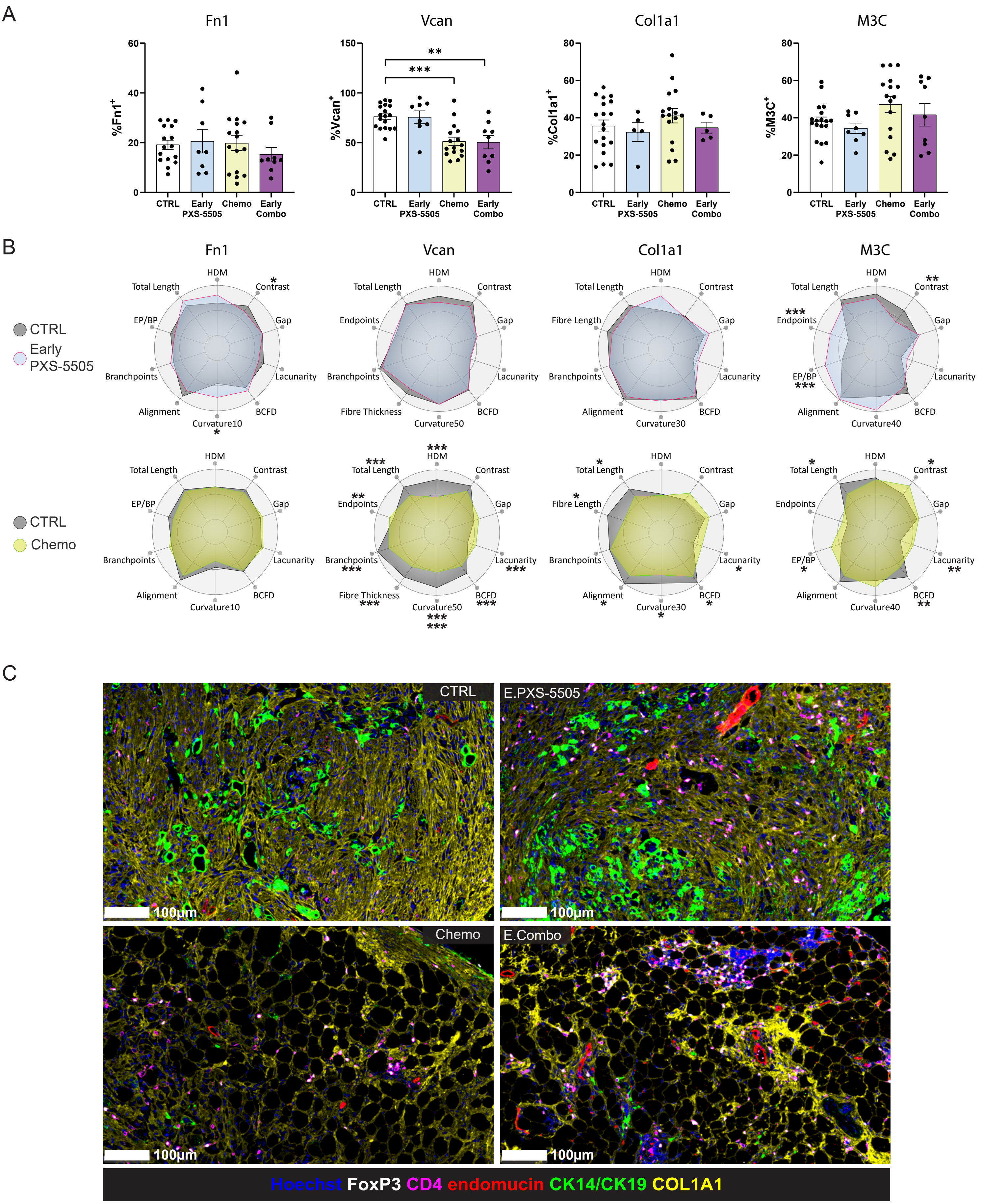

### Supplemental Figure 6

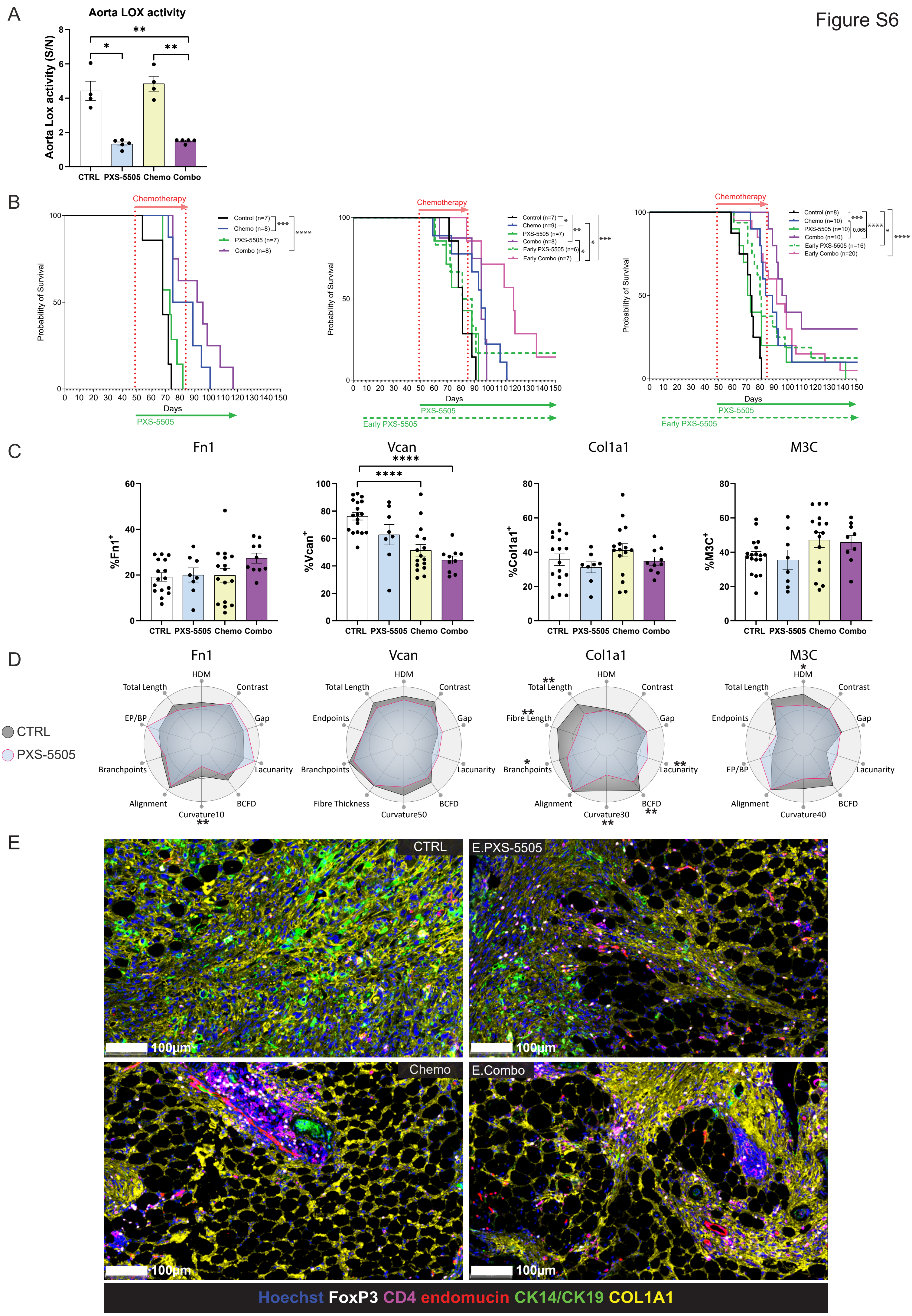
